# The β6 Integrin Negatively Regulates TLR7-Mediated Epithelial Immunity via Autophagy During Influenza A Virus Infection

**DOI:** 10.1101/2023.08.28.555098

**Authors:** Maria Smith, Victoria Meliopoulos, Shaoyuan Tan, Theresa Bub, Pamela H. Brigleb, Bridgett Sharp, Jeremy Chase Crawford, Mollie S. Prater, Shondra M. Pruett-Miller, Stacey Schultz-Cherry

## Abstract

Integrins are essential surface receptors that sense extracellular changes to initiate various intracellular signaling cascades. The rapid activation of the epithelial-intrinsic β6 integrin during influenza A virus (IAV) infection has been linked to innate immune impairments. Yet, how β6 regulates epithelial immunity remains undefined. Here, we identify the role of β6 in mediating the Toll-like receptor 7 (TLR7) through the regulation of intracellular trafficking. We demonstrate that deletion of the β6 integrin in lung epithelial cells significantly enhances the TLR7-mediated activation of the type I interferon (IFN) response during homeostasis and respiratory infection. IAV-induced β6 facilitates TLR7 trafficking to lysosome-associated membrane protein (LAMP2a) components, leading to a reduction in endosomal compartments and associated TLR7 signaling. Our findings reveal an unappreciated role of β6-induced autophagy in influencing epithelial immune responses during influenza virus infection.

## INTRODUCTION

The respiratory epithelium serves as a critical interface between the inhaled environment and the host microenvironment (Denney and Ho, 2018). Due to its unique position, this epithelial barrier can respond to stimuli from the external environment and signal to the immune system to protect the underlying mesenchyme (Parker and Prince, 2011). As a result, the lung epithelium plays an indispensable role in maintaining tissue homeostasis by facilitating the mucociliary clearance of noxious pathogens and conferring frontline defenses via its antigen-sensing and immunomodulatory mechanisms (Kim *et al*., 1997; Whitsett and Alenghat, 2014; Iwasaki, Foxman and Molony, 2017; Kuek and Lee, 2020). However, inhaled pathogens can disrupt barrier integrity by increasing cellular permeability, deterring epithelial cell regeneration, and dampening antiviral immune responses (Couceiro, Paulson and Baum, 1993; Short *et al*., 2014; Meliopoulos *et al*., 2016). A fundamental question that remains is how epithelial-intrinsic surface receptors modulate innate immunity during viral infection.

Integrins are heterodimeric cell surface receptors that interact with the extracellular matrix and associate with the cytoskeleton to initiate intracellular signaling pathways (Hood and Cheresh, 2002; Anderson, Owens and Naylor, 2013). These heterodimers consist of ⍺ and β subunits, which regulate numerous processes including cell adhesion and activation, tissue morphogenesis, and cytoskeletal remodeling (Sheppard, 2003). The β6 integrin (*Itgβ6*) preferentially associates with the ⍺v subunit, resulting in low expression of the ⍺vβ6 heterodimer exclusively in epithelial cells, where it plays a critical role in the maintenance of lung tissue homeostasis and activation of anti-inflammatory mechanisms (Breuss *et al*., 1993, 1995; Huang and Farese, 1996). However, the β6 integrin is also implicated in multiple models of lung fibrosis and injury due to its rapid activation during epithelial injury, inflammation, and viral infections such as influenza virus (Huang and Farese, 1996; J. S. Munger *et al*., 1999; Sheppard, 2003; Horan *et al*., 2008; Puthawala *et al*., 2008; Meliopoulos *et al*., 2016; Decaris *et al*., 2021).

In conjunction with integrins, pathogen recognition receptors called Toll-like receptors (TLRs) also play an essential role in regulating host innate immunity (Parker and Prince, 2011; Iwasaki and Pillai, 2014; Whitsett and Alenghat, 2014). During influenza A virus (IAV) infection, TLRs are responsible for transducing antiviral signaling cascades that sequentially activate the Nuclear Factor-кB 1 (NFкB1) and the Interferon (IFN) Regulatory Factors (IRFs) to initiate robust antiviral responses that interfere with viral replication, recruit immune cells to the site of infection, and facilitate viral clearance (Sato *et al*., 2003; Moynagh, 2005; Kawai and Akira, 2007; Ioannidis *et al*., 2013). In particular, the endosomal-localized TLR7 sensor is involved in the recognition and elimination of single-stranded RNA by inducing pro-inflammatory cytokines and type I IFNs against influenza virus and many other microbial infections (Testerman *et al*., 1995; Diebold *et al*., 2004; Ioannidis *et al*., 2013; Iwasaki and Pillai, 2014; Fan, Ren and Hou, 2018; van der Sluis *et al*., 2022). Cell-extrinsic recognition of antigens is also critically mediated by TLR7 through the recruitment of innate immune cells, activation of antigen-presenting cells, and induction of adaptive immunity (Koyama *et al*., 2007; Hou, Reizis and DeFranco, 2008; Di Domizio *et al*., 2009; Kang *et al*., 2011; Jeisy-Scott *et al*., 2012). Despite the significance of TLR7-mediated signaling in orchestrating robust antiviral defenses against pathogens, the mechanisms by which TLR7 signaling is regulated remain understudied.

TLR7 activation is dependent on its subcellular localization within endosomal compartments (Petes, Odoardi and Gee, 2017). However, these endosomes can be sequestered by intracellular compartments derived from the autophagy pathway (Gordon and Seglen, 1988; Ganesan and Cai, 2021). Macroautophagy is a catabolic process induced during nutrient deprivation and cellular stress for the degradation of damaged organelles and intracellular pathogens (Parzych and Klionsky, 2014; Omotade and Roy, 2019). This pathway involves the formation of double-membrane autophagosome vesicles that subsequently fuse with acidic lysosomes to facilitate the degradation of engulfed cytoplasmic contents (Parzych and Klionsky, 2014). Although IAVs have developed strategies to manipulate autophagy machinery to counteract antiviral responses, our understanding of the mechanisms by which IAV-induced autophagy modulates TLR7 signaling remains limited (Perot *et al*., 2018; Wang *et al*., 2019, 2020; Zhou *et al*., no date).

Our previous studies have shown that seasonal IAV significantly upregulates the β6 integrin early during infection, which increased host susceptibility, disease pathogenesis, and mortality (Meliopoulos *et al*., 2016). The absence of the β6 integrin constitutively activated type I IFN signaling that significantly reduced viral spread, minimized lung injury, and considerably improved host resistance and survival (Meliopoulos *et al*., 2016). This protective phenotype extended beyond this seasonal IAV strain, as β6KO mice were also resistant to avian IAV, Sendai virus, and *Streptococcus pneumonia* infections (Meliopoulos *et al*., 2016). While our study identified the suppressive role of β6 on lung-intrinsic immune mechanisms, additional studies are warranted to delineate the mechanisms by which IAV-induced β6 negatively regulates epithelial-derived type I IFN responses.

Here, we identify an epithelial-intrinsic role of the β6 integrin in restricting TLR7 signaling. The absence of the β6 integrin significantly upregulated the type I IFN response via a TLR7-dependent mechanism, resulting in elevated NFкB1 and STAT1/p-STAT1 signaling. Mechanistically, we demonstrated that IAV-induced β6 promotes the recruitment of autophagy machinery in lung epithelial cells. This led to significant colocalization between TLR7 and LAMP2a lysosomal compartments and reduced endosome availability, which subsequently inhibited TLR7 activation and signaling. These studies reveal a previously unrecognized association between the β6 integrin and autophagy machinery as an epithelial-intrinsic mechanism of regulating TLR7 signaling and epithelial immunity during influenza A virus infection.

## RESULTS

### *Itg*β*6* knockout enhances TLR7 and NFкB1 signaling at baseline

Our previous studies demonstrated a protective phenotype in β6-deficient (β6KO) mice (Meliopoulos *et al*., 2016). Although the increased resistance against IAV challenge has been attributed to a type I IFN-dependent mechanism, the specific IFN factors targeted by β6 to modulate this antiviral response remain unknown. Therefore, we hypothesized that the β6-mediated regulation of the type I IFN response occurs through the modulation of upstream IFN intermediates. To test this, primary murine tracheobronchial epithelial cells (mTECs) were isolated from C57Bl/6 wildtype (WT) and β6KO mice and differentiated at the air-liquid interface (Meliopoulos *et al*., 2016). Among the endosomal TLRs, RNA-sequencing (RNA-seq) analyses revealed the highest upregulation of *Tlr7* in β6KO mTECs at baseline, including significant elevation of *Ifih1*, *Casp1*, and *Nfκb1* (Figure 1A). Similarly, a murine type I IFN RT^2^ profiler array revealed upregulation of *Tlr7*, *Tlr9*, *Myd88*, *Irfs*, and Interferon Stimulated Genes (ISGs) in β6KO mTECs at baseline compared to WT mTECs (Figure 1B). In accordance with these transcriptional analyses, confocal microscopy also revealed significantly higher TLR7 expression in uninfected β6KO mTECs (Figure 1C). Following TLR7 activation, signals are subsequently transduced to NFκB1 to prompt IFN release (Moynagh, 2005; Kawai and Akira, 2007). Therefore, WT and β6KO mTECs were stained for the p65 subunit of NFκB1 (Anrather *et al*., 1999). β6KO mTECs exhibited substantially more NFкB1 expression, indicative of increased TLR7 signaling (Figure 1D).

**Figure 1.**
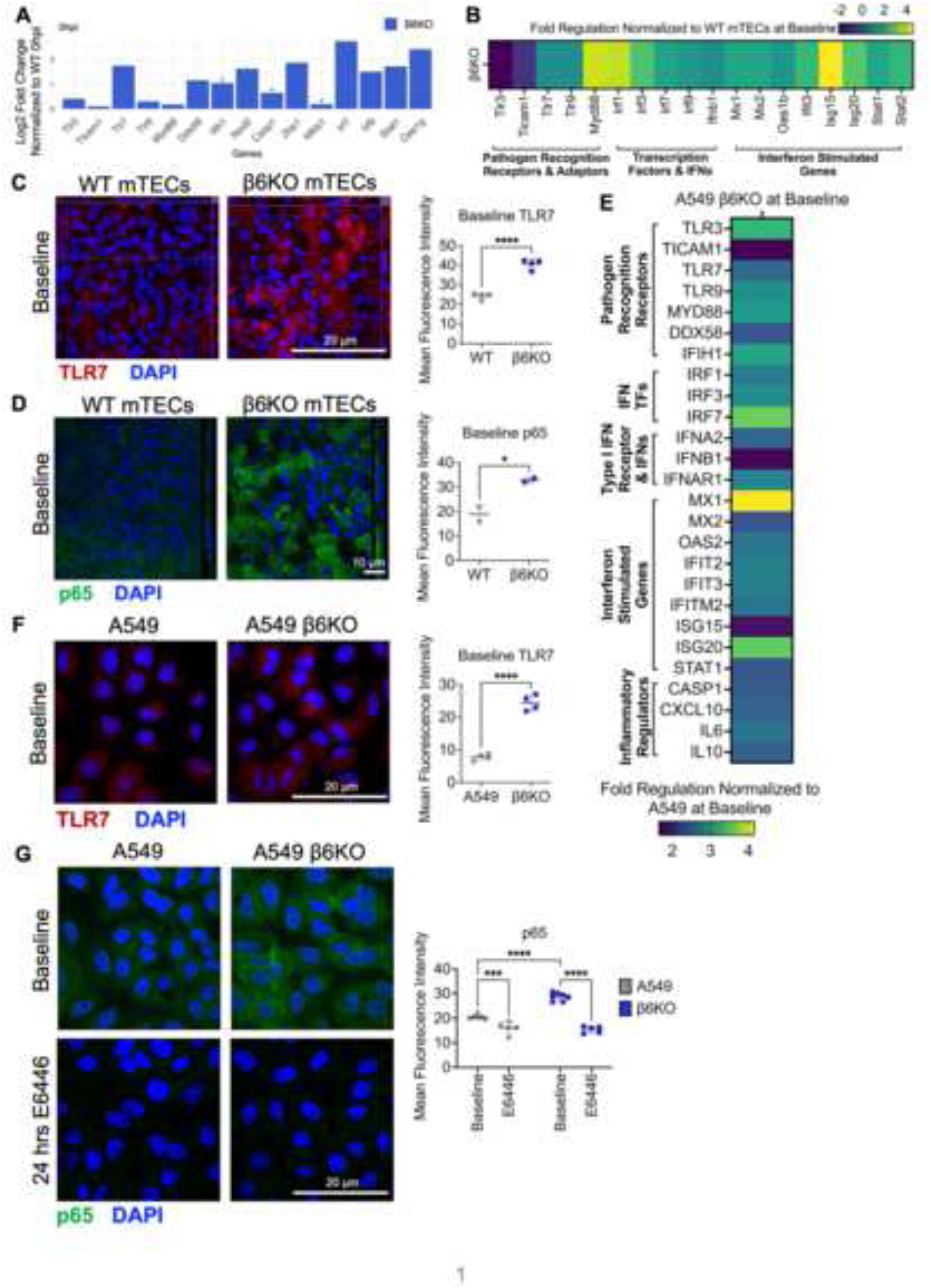
Itgβ6 knockout enhances TLR7 and NFкB1 signaling at baseline. (A) RNA-sequencing analysis of type I interferon-regulated genes from differentiated WT and β6KO mTECs at baseline from two independent experiments. Data reflective of expression in β6KO mTECs (represented in blue) relative to WT mTECs. Dataset from (Meliopoulos *et al*., 2016). *p≤0.05. (B) Heatmap depicting fold-regulation of the type I interferon response in β6KO mTECs at baseline relative to uninfected WT mTECs. Data analyzed with a murine type I interferon RT^2^ Profiler from 2 independent experiments with n=2 mTECs pooled per group. Yellow indicates higher expression and purple indicates lower expression. (C) Uninfected WT and β6KO mTECs were immunostained for TLR7 (red) and DAPI (blue). Data is pooled from 2 independent experiments with n=2 mTECs per group. Scale bar, 20µm. (D) Uninfected WT and β6KO mTECs were immunostained for NFκB subunit p65 (green) and nucleus (Hoechst, blue). Data is representative of 2 independent experiments with n=2 mTECs per group. Scale bar, 10µm. (E) Heatmap depicting fold-regulation of the type I interferon response in A549 β6KO cells at baseline relative to uninfected A549 cells. Data were analyzed with a human type I interferon RT^2^ Profiler from 3 independent experiments with n=2 cells per group. Yellow indicates higher expression and purple indicates lower expression. TFs = Transcription Factors. (F) Uninfected A549 and A549 β6KO cells were immunostained for TLR7 (red) and nucleus (Hoechst, blue). Data is pooled from 2 independent experiments with n=2 cells imaged per group. Scale bar, 20µm. (G) Uninfected A549 and A549 β6KO cells were immunostained for NFκB subunit p65 (green), and nucleus (Hoechst, blue). Top panel indicates baseline samples while bottom panel indicates uninfected samples that were treated with 10µM E6446 TLR7 inhibitor for 24 hours. Data is pooled from 2 independent experiments with n=2-3 cells imaged per group. Scale bar, 20µm. Mean fluorescence intensity quantified using Fiji. Data graphed as mean ± SD via unpaired T-test. *p<0.05; ***p≤0.0001; ****p<0.0001. See also Figure S1.

Next, we investigated whether this phenotype could be recapitulated using human lung epithelial cells. Within the respiratory epithelium, alveolar epithelial cells are critical for the preservation of lung homeostasis and the activation of innate immune responses (Guillot *et al*., 2013). Therefore, we utilized the adenocarcinoma human alveolar basal epithelial (A549) cell line that has been widely used to investigate the impact of viral infections on the lung epithelium (Atkins *et al*., 2014). Notably, A549s also express high endogenous β6 levels (Yan *et al*., 2018). Therefore, we generated an A549 β6KO cell line via CRISPR-Cas9 knockout of the *Itgβ6* gene. Relative to the A549 β6KO cell line, β6 expression was significantly upregulated in A549 cells at baseline (Figures S1A and S1B). In alignment with the IFN profile observed in β6KO whole lungs and mTECs, heightened transcripts associated with type I IFN-mediated recognition, signaling, and induction were evident in uninfected A549 β6KO cells (Figure 1E). Confocal microscopy also demonstrated a substantial increase of TLR7 and NFкB1 in A549 β6KO cells, compared to A549 cells (Figures 1F and 1G). Subsequently, this led us to investigate whether the enhanced immune response in β6KO epithelial cells was driven by a TLR7-dependent mechanism. To test this, we treated A549 and A549 β6KO cells with the E6446 TLR7 antagonist for 24 hours (Lamphier *et al*., 2014). E6446 inhibits TLR7 binding and signaling by interfering with the TLR7 receptor domain, which destabilizes the association between the viral sensor and nucleic acids (Figure S1C) (Lamphier *et al*., 2014). E6446 treatment led to blunting of the NFкB1 response not only in the A549 cells but also in the A549 β6KO cells (Figure 1G). These data suggest the physiological role of β6 in regulating epithelial-derived IFN responses via modulation of TLR7 and NFкB1.

### Influenza-induced β6 suppresses TLR7 signaling to mediate the type I IFN response

To investigate the role of β6 in modulating the innate immune response during microbial insults, we infected A549 and A549 β6KO cells with A/California/04/2009 (CA/09) H1N1 virus at a multiplicity of infection (MOI) 1 for 24 hours post-infection (hpi). In response to infection, A549 β6KO cells produced higher STAT1 levels and significantly more phosphorylated STAT1 (pSTAT1) at 24 hpi, indicative of increased type I IFN induction (Figure 2A). Given the upregulation of TLR7 and NFкB1 in the β6KO model at baseline, we assessed whether this antiviral phenotype was maintained upon IAV infection. Confocal microscopy continued to indicate significantly higher fluorescence of TLR7 and NFкB1 in the A549 β6KO cells at 24 hpi, to a greater extent than observed in infected A549 cells (Figures 2B and 2C). Similarly, CA/09-infected β6KO mTECs expressed considerably higher TLR7 and NFкB1 expression at 8 hpi and 24 hpi (Figure S2A and S2B). Although WT mTECs mounted a TLR7 response against infection, NFкB1 expression was delayed until 24 hpi (Figure S2B).

**Figure 2.**
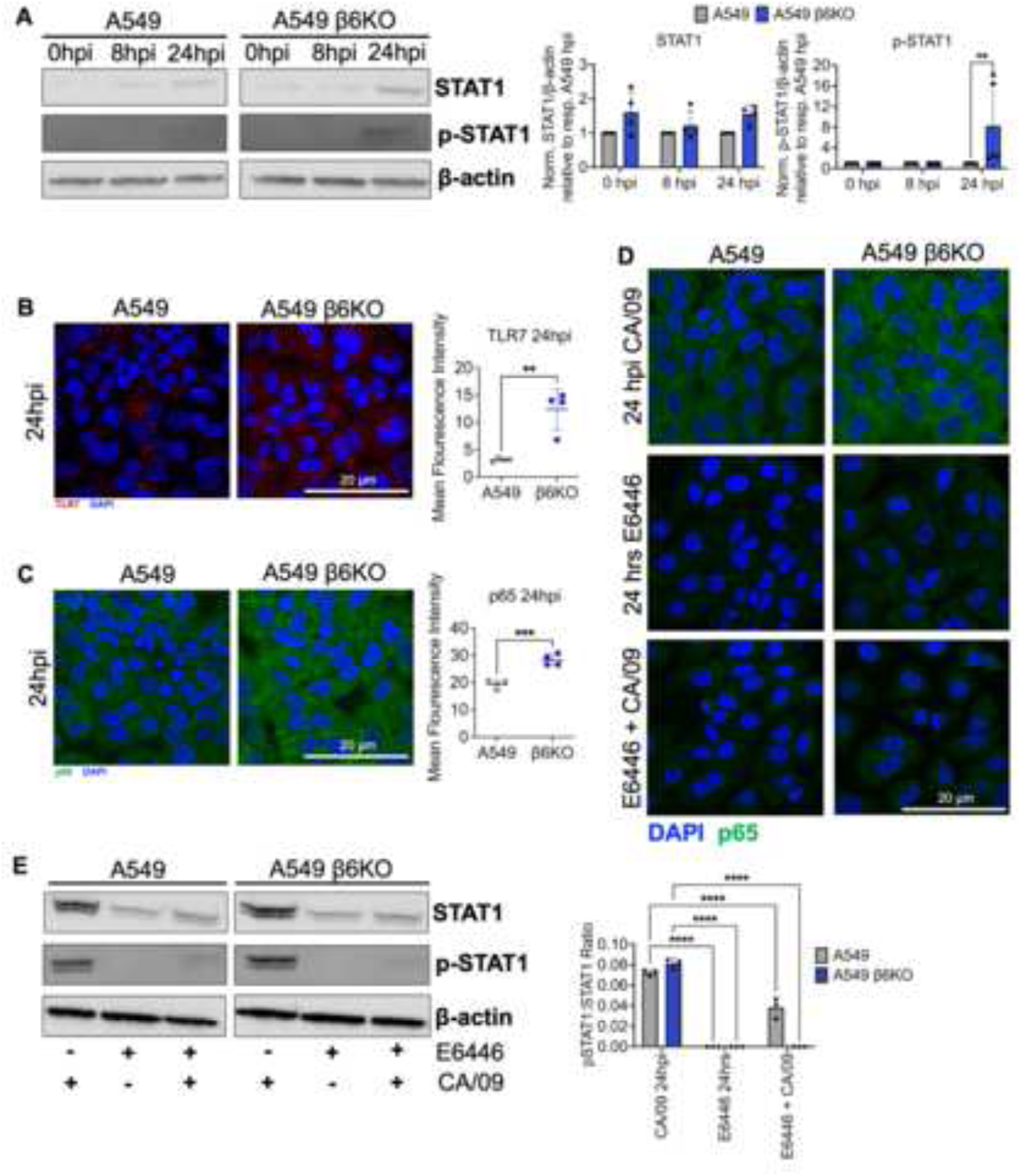
Influenza-induced β6 suppresses TLR7 signaling to mediate the type I IFN response. (A) Immunoblot analysis of STAT1, phosphorylated STAT1 (p-STAT1), and β-actin in the lysates of A549 and A549 β6KO cells at 0, 8, and 24 hpi. Infected cells were incubated with CA/09 virus MOI 1 for 8 or 24 hours. Data is pooled from 2 independent experiments with n=2-3 cells per group. Normalized STAT1 and p-STAT1 from A549 β6KO cells were normalized to A549 cells at each respective time point. (B) A549 and A549 β6KO cells were infected with CA/09 virus MOI 1 for 24 hours and immunostained for TLR7 (red) and nucleus (Hoechst, blue). Data is pooled from 2 independent experiments with n=2 cells per group. Scale bar, 20µm. (C) A549 and A549 β6KO cells were infected with CA/09 virus MOI 1 for 24 hours and immunostained for NFκB subunit p65 (green) and nucleus (Hoechst, blue). Data is pooled from 2 independent experiments with n=2 cells per group. Scale bar, 20µm. (D) A549 and A549 β6KO cells were immunostained for NFκB subunit p65 (green) and nucleus (Hoechst, blue). Data is representative of 2 independent experiments with n=3 cells per group. First panel indicates cells that were infected with CA/09 virus for 24 hpi. Second panel consists of cells treated with 10µM E6446 TLR7 inhibitor for 24 hours. Third panel indicates cells pre-treated with 10 µM E6446 and then infected with CA/09 virus MOI 1 for 24 hours. Scale bar, 20µm. (E) Immunoblot analysis of STAT1, phosphorylated STAT1 (p-STAT1), and β-actin in the lysates of A549 and A549 β6KO cells treated as described in D: first column=CA/09 virus only; second column=E6446 only; third column=E6446 + CA/09. Mean fluorescence intensity quantified using Fiji version. Data graphed as mean ± SD. Two-way ANOVA with Sidak’s multiple comparisons test for A and E; Unpaired T-test for B and C. **p<0.005; ***p≤0.0001; ****p<0.0001. See also Figures S1 and S2.

Additionally, we evaluated the dependence on TLR7 in initiating and driving the antiviral response in β6KO cells. To test this, A549 and A549 β6KO cells were either infected with CA/09 virus at MOI 1 for 24 hpi, treated with E6446 for 24 hours, or pre-treated with E6446 for 24 hours and then infected with CA/09 virus for 24 hpi (E6446 + CA/09). While CA/09 infection induced NFкB1 expression in both cell lines, E6446 treatment attenuated NFкB1 expression (Figure 2D). E6446 + CA/09 treatment also resulted in significant downregulation of NFкB1 expression, hindering the ability of epithelial cells to mount TLR7-mediated NFкB1 responses against IAV infection (Figure 2D). Furthermore, we assessed whether TLR7 signaling facilitated IFN induction in β6KO cells. At 24 hpi, CA/09 virus stimulated STAT1 and pSTAT1 upregulation in both cell lines as anticipated (Figure 3E). In comparison, E6446 treatment led to a significant loss of STAT1 and a complete loss of pSTAT1 (Figure 3E). E6446 + CA/09 treatments induced a similar inhibitory effect on the STAT1/pSTAT1 response in A549 and A549 β6KO cells, highlighting that TLR7 is crucial for driving the enhanced antiviral response in β6KO cells. Together, these data indicate that IAV-induced β6 can exert an inhibitory effect on the TLR7-driven type I IFN response in respiratory epithelial cells.

**Figure 3.**
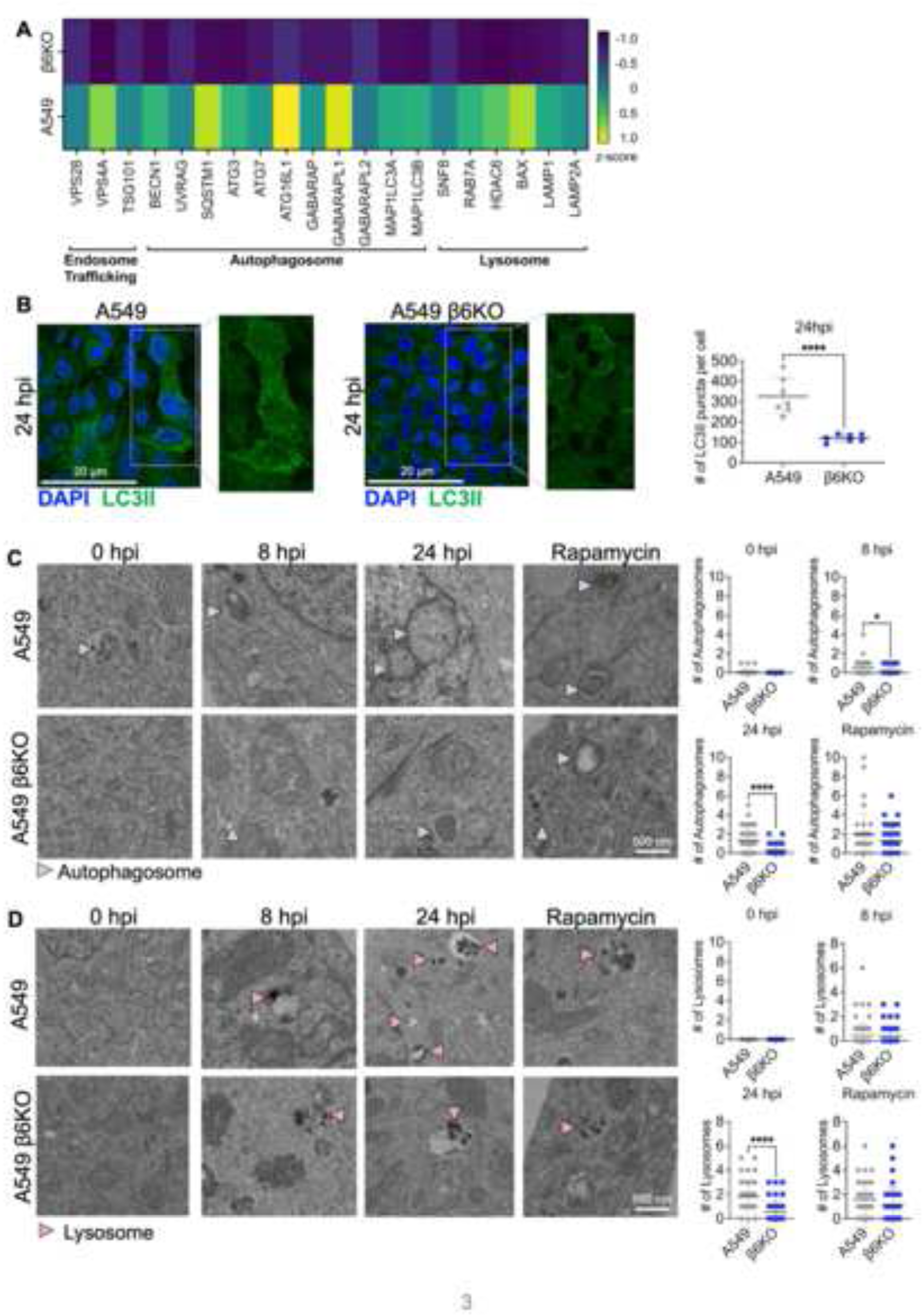
The β6 integrin increases autophagy induction during influenza infection. (A) Heatmap comparing the expression of endosomal-, autophagy- and lysosomal-associated genes between A549 and A549 β6KO cells at 8 hours post-CA/09 virus infection (MOI 1). Pooled samples from 2 independent experiments with n=3 cells per group were analyzed with a custom autophagy RT^2^ Profiler by z-score. Yellow indicates higher expression and purple indicates lower expression. (B) A549 and A549 β6KO cells were infected with CA/09 virus MOI 1 for 24 hours and immunostained for LC3-II (green), and nucleus (Hoechst, blue). Data is pooled from 2 independent experiments with n=3-4 cells per group. Number of LC3-II puncta per cell was quantified using Fiji. Scale bar, 20µm. (C-D) Transmission electron microscopy (TEM) of autophagy machinery in A549 and A549 β6KO cells at baseline, 8 and 24 hpi CA/09 MOI 1 infection and 24-hour 100nM rapamycin treatment. Data were pooled from 2 independent experiments with n=1 cell per group for each treatment condition and quantified from 63-120 unbiased images for autophagosome and lysosome quantifications. (C) Blue arrows indicate double-membrane autophagosomes. (D) Orange arrows indicate lysosomes. Scale bar, 500nm. Data graphed as mean ± SD via unpaired T-test. *p<0.05; ****p<0.0001. See also Figure S3.

### The β6 integrin increases autophagy induction during influenza infection

Given the sequestration of TLR7 viral sensors within endosomes, we hypothesized that the β6-mediated suppression of TLR7-driven immunity occurred via an intracellular mechanism capable of regulating endosomal vesicles. A recent study reported that the regulation of TLR signaling occurred via the integrin-mediated fusion of TLR-containing endosomes to the lysosome, which induced endosomal degradation and attenuated TLR activity (Acharya *et al*., 2016). While studies have observed autophagy induction in IAV-infected respiratory epithelial cells, none have attributed the β6-mediated trafficking of endosomes to blunted TLR signaling during IAV infection. First, to assess if β6 upregulated autophagy induction during IAV infection, we utilized a customized autophagy Reverse Transcription-Polymerase Chain Reaction (RT-PCR) array that included genes related to endosomal trafficking, autophagosome formation along with lysosomal fusion and degradation. At 8hpi, IAV-infected A549 cells indicated significant upregulation of processes related to endosomal transport, autophagosome maturation, autolysosome formation, and lysosomal-mediated degradation, compared to A549 β6KO cells (Figure 3A). Confocal microscopy also showed increased autophagy induction, as evidenced by prominent microtubule-associated protein light chain 3-II (LC3-II) puncta that were observed to a significantly higher extent in CA/09-infected A549 cells (Figure 3B) (Tanida, Ueno and Kominami, 2004).

To further confirm this phenotype, we performed transmission electron microscopy (TEM) on A549 and A549 β6KO cells that were either uninfected, infected with CA/09 virus MOI 1 for 8 hpi and 24 hpi or treated for 24 hours rapamycin, an autophagy inducer (Ravikumar *et al*., 2004). While a few autophagosomes were detected exclusively in the A549s at baseline, a more distinctive phenotype was observed upon IAV infection (Figure 3C). The formation of double-membrane autophagosomes was prevalent in A549 cells at 8 hpi and 24 hpi, in comparison to A549 β6KO cells where substantially fewer autophagosomes were identified (Figure 3C). Similarly, IAV infection recruited significantly more lysosomes in 24-hour infected A549 cells to higher levels than detected in the infected A549 β6KO cells (Figure 3D). In addition to this, amphisomes and autolysosomes were exclusively detected in 24-hour infected A549 cells, indicating activation of endosome-autophagosome and autophagosome-lysosome fusion mechanisms, respectively (Figure S3A and S3B). Altogether, these findings indicate that the β6 integrin is critical for increasing the recruitment of autophagy machinery during IAV infection.

### The β6 integrin promotes the trafficking of TLR7 to the lysosome

The intracellular trafficking and localization of TLR7 within the endosomal compartment are crucial for its activation, antigen-sensing potential, and signaling capabilities (Petes, Odoardi and Gee, 2017). Since integrins influence subcellular mechanisms via their cytoplasmic tail domains, we hypothesized that the β6-mediated modulation of epithelial immunity could be attributed to effects on TLR7 intracellular trafficking (Zhang and Wang, 2012). To test this, A549 and A549 β6KO cells were infected with CA/09 virus, and then confocal microscopy was used to identify the intracellular localization of TLR7. In A549 cells, we observed a significant reduction in early endosomes (EEA1) at 8 and 24 hpi, which was accompanied by a substantial loss of TLR7 signal (Figure 4A and S4A). Although substantially more endosomes and higher TLR7 expression were identified in A549 β6KO cells, co-localization between TLR7 and EEA1 was not observed at 24hpi (Figure 4A and S4A).

**Figure 4.**
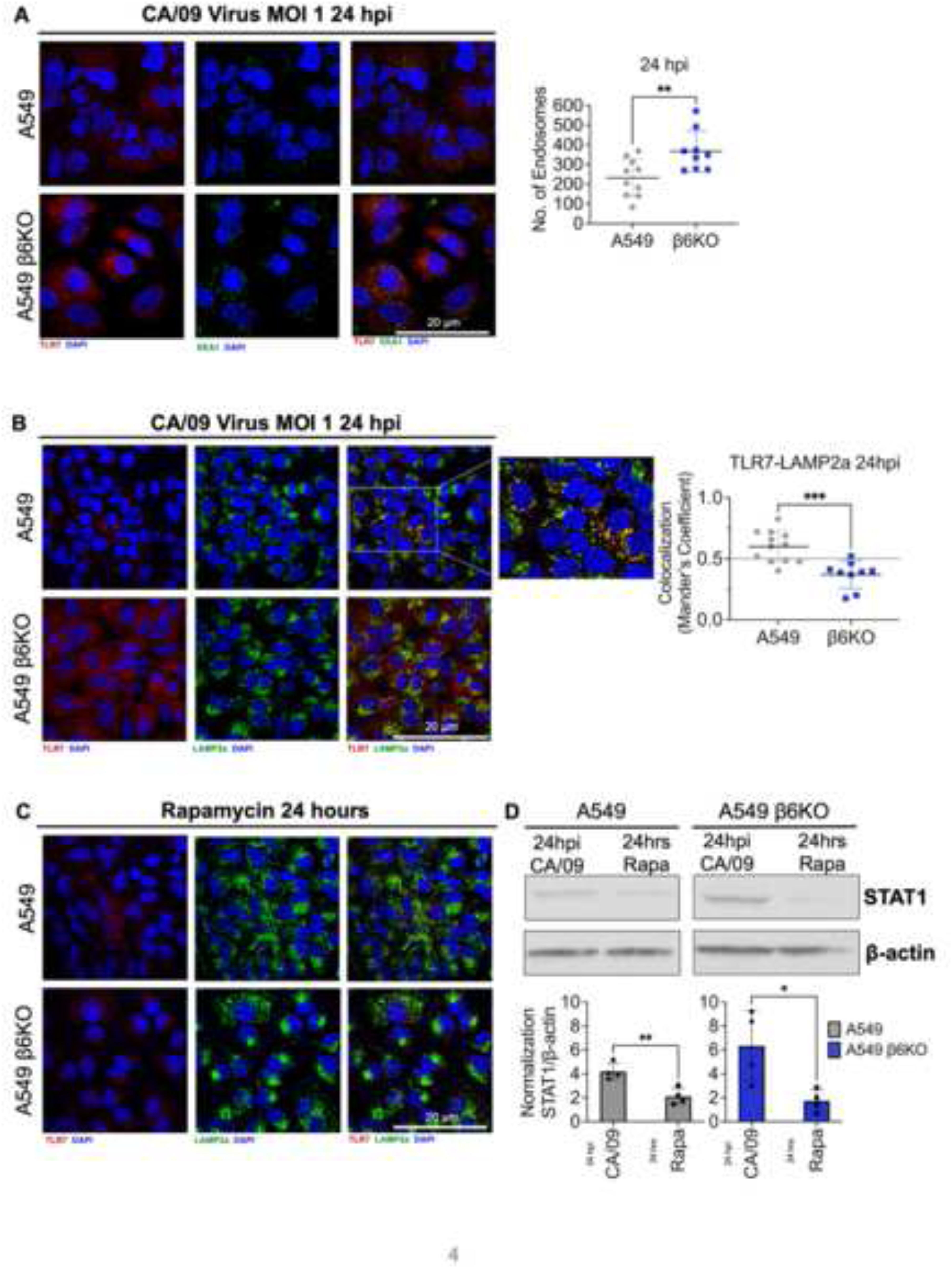
The β6 integrin promotes the trafficking of TLR7 to the lysosome. (A) A549 and A549 β6KO cells were infected with CA/09 virus MOI 1 for 24 hours and immunostained for TLR7 (red), EEA1 endosome (green), and nucleus (Hoechst, blue). Data is pooled from 2 independent experiments with n=4-5 cells per group and the number of endosomes was quantified with Fiji. Scale bar, 20µm. (B) A549 and A549 β6KO cells were infected with CA/09 virus MOI 1 for 24 hours and immunostained for TLR7 (red), LAMP2a lysosome (green), and nucleus (Hoechst, blue). TLR7-LAMP2a co-localization quantified with Mander’s co-efficient (>0.5=colocalized; <0.5=not colocalized). Data were pooled from 2 independent experiments with n=4-6 cells imaged per group. Scale bar, 20µm. (C) A549 and A549 β6KO cells were incubated with 100nM rapamycin for 24 hours and immunostained for TLR7 (red), LAMP2a lysosome (green), and nucleus (Hoechst, blue). Data representative of 2 independent experiments with n=2-3 cells imaged per group. Scale bar, 20µm. (D) Immunoblot analysis of STAT1 and β-actin in the lysates of A549 and A549 β6KO cells infected with CA/09 virus MOI 1 or treated with 100nM rapamycin for 24 hours. Data representative of 2 independent experiments with n=4 cells per group. Data graphed as mean ± SD via unpaired T-test. *p<0.05; **p<0.005; ***p≤0.0001. See also Figure S4.

Considering that we detected intermediate amphisome and autolysosomal organelles in A549 cells at 24 hpi, we hypothesized that β6 facilitated associations between TLR7 and terminal lysosomal machinery that led to a substantial loss of endosomes in β6-expressing cells (Figures S3A and S3B). Following CA/09 infection, TLR7 significantly colocalized with the lysosome-associated membrane protein (LAMP2a) in A549 cells (Mander’s coefficient >0.5), suggestive of trafficking to the lysosomes at 24 hpi (Figure 4B) (Dunn, Kamocka and McDonald, 2011). However, TLR7 did not colocalize with the lysosomal compartments in infected A549 β6KO cells (Mander’s coefficient <0.5), demonstrating that β6 is essential for mediating TLR7 trafficking (Figure 4B).

Lastly, we determined whether β6 regulation of epithelial-derived immunity was specific to IAV infection. To explore this, we induced autophagy in both cell lines through 24-hour rapamycin treatment, which significantly reduced the TLR7 response in both A549 and A549 β6KO cells (Figure 4C). Notably, recruitment of LAMP2a was observed to a higher extent in the A549s than in the A549 β6KO cells (Figure 4C). This observation further supports the identified role of β6 in increasing the induction of autophagy machinery (Figure 3D). In comparison to infected cells, rapamycin-treated cells presented a significant decrease in STAT1 signal, suggesting a broader role of autophagy in regulating epithelial immune mechanisms beyond the scope of viral infections (Figure 4D). These data confirm that the β6 integrin is critical for the TLR7 intracellular relocalization process, as the formation of LC3-puncta, detection of intermediary autophagy machinery, and co-localization of TLR7 with LAMP2 were significantly reduced in β6-deficient epithelial cells. They also indicate that autophagy-mediated regulation of innate immunity can occur independent of IAV infection. Together, these data propose a mechanism whereby β6 modulates epithelial immunity through the increased recruitment of terminal autophagy machinery and their association with the TLR7 viral sensor.

### Single-cell RNA sequencing indicates that β6-positive lung epithelial cells from wildtype mice exhibit an attenuated type I IFN response and upregulated autophagy induction during IAV infection

To determine the relevance of the β6 integrin in regulating epithelial immunity in an *in vivo* model, we utilized the single-cell RNA-sequencing (scRNA-seq) dataset courtesy of Dr. Paul Thomas (Boyd *et al*., 2020). Briefly, whole lungs were isolated from C57Bl/6 wildtype mice that were uninfected or infected with influenza A/Puerto Rico/8/34 (PR8) virus, which is a lethal mouse-adapted IAV strain commonly used to investigate host immunity and pathogenesis during IAV infections (Figure 5A) (Rutigliano *et al*., 2014; Boyd *et al*., 2020). Lung single-cell suspensions were sorted for CD45-negative cells, allowing the identification of distinct cell populations that consisted primarily of endothelial, mesenchymal, and epithelial cells (Figure 5B). Among the epithelial population, we identified club, ciliated, type I alveolar (ATI), and type II alveolar (ATII) epithelial cells (Figure 5B). Expression analyses revealed significantly blunted type I IFN expression in β6-positive lung epithelial cells at baseline and 1-day post-infection (dpi) (Figures 5C, 5D, 5E, 5F). To further examine this blunted IFN response, we assessed potential IFN targets that might be subject to β6 regulation. Consistent with the downregulated IFN targets in A549 cells, *Myd88*, *Nfкb1*, *Ifn⍺r1*, *Ifn⍺r2*, and *Stat1* levels were significantly reduced in β6-positive lung epithelial cells at baseline, which remained suppressed at 1 dpi and 3 dpi (Figure 5G). Finally, we compared the expression of autophagy- and lysosomal-related genes between infected β6-positive and β6-negative epithelial cells. At 24 hpi, *Atg3*, *Atg5*, *Atg7*, *Atg12*, *Gabarapl1*, *Gabarapl2*, *Map1lc3b*, *Rab7*, *Lamp1* and *Lamp2* were upregulated in β6-positive cells but not in the β6-negative cells (Figure 5H). Taken together, this dataset provides new evidence supporting the potential role of β6 in shaping the epithelial responses against various IAV strains through its influence on IFN and autophagy pathways.

**Figure 5.**
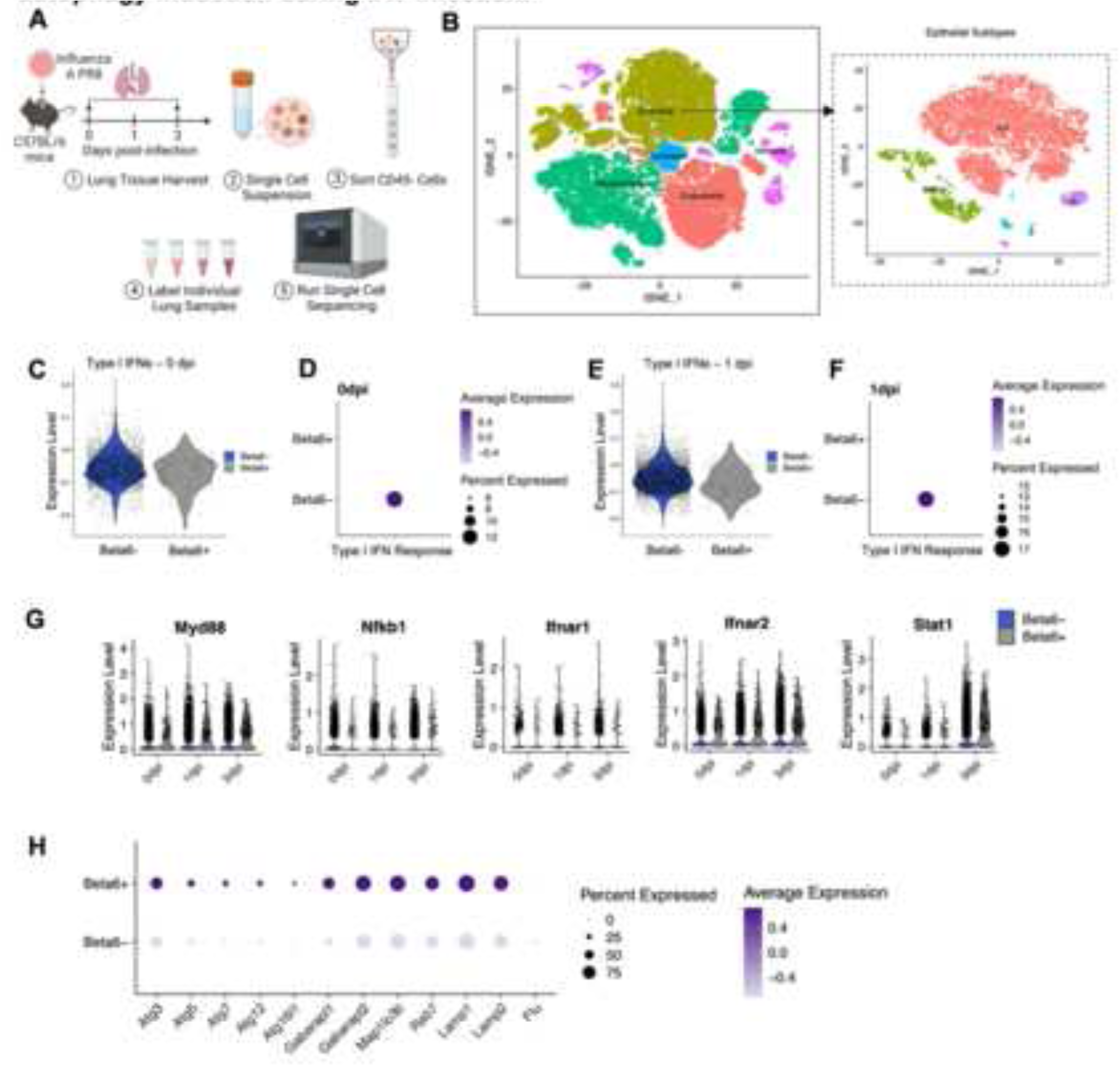
Single-cell RNA sequencing indicates that β6-positive lung epithelial cells from wildtype mice exhibit an attenuated type I IFN response and upregulated autophagy induction during IAV infection. (A) Schematic of scRNA-sequencing process, adapted from (Boyd *et al*., 2020). Whole lungs were harvested from uninfected and infected (IAV/Puerto Rico/8/34 virus) WT mice at 0-, 1-and 3-dpi. CD45-negative cells were sorted from single-cell lung suspensions, individually labeled, and sequenced. (B) *t*-distributed stochastic neighbor embedding (*t*-SNE) projection of murine lung cell populations and identified lung epithelial subsets. (C-F) Violin and dot plots of the type I interferon response in the total lung epithelial cell population. The comprehensive type I interferon signature included the following genes: *Nod1*, *Nod2*, *Casp1*, *Nlrp3*, *Aim2*, *Tlr3*, *Ticam1*, *Tlr7*, *Tlr9*, *Myd88*, *Ddx41*, *Trim56*, *Zbp1*, *Ripk3*, *Nfkb1*, *Irf7*, *Irf9*, *Tyk2* and *Stat1*. Type I IFN expression compared between β6-negative (β6-) and β6-positive (β6+) epithelial cells at (C-D) 0 dpi and (E-F) 1 dpi. (G) Expression of type I IFN-mediators, *Myd88*, *Nfkb1*, *Ifnɑr1*, *Ifnɑr2*, *Stat1*, between β6- and β6+ epithelial cells at 0, 1 and 3 dpi PR8 inoculation. (H) Dot plot of autophagy- and lysosome-associated genes in β6-versus β6+ epithelial cells at 1 dpi PR8 inoculation. Data are mean ± SEM.

## DISCUSSION

Our data reveal a new mechanism by which the β6 integrin mediates epithelial IFN responses through the modulation of TLR7 signaling. We demonstrate that IAV-induced β6 promotes the recruitment of autophagy machinery in alveolar epithelial cells that facilitate increased trafficking of TLR7 sensors to the lysosomes, leading to a significant reduction in endosome availability and type I IFN signaling (Figure 6A). The absence of the β6 integrin decreases autophagy induction, which increases endosome availability and considerably improves TLR7-mediated NFкB1 signaling and the type I IFN response against IAV infection (Figure 6B). Consequently, we propose the role of an epithelial-intrinsic negative regulator that counteracts innate viral sensing and IFN signaling mechanisms to influence host susceptibility.

**Figure 6.**
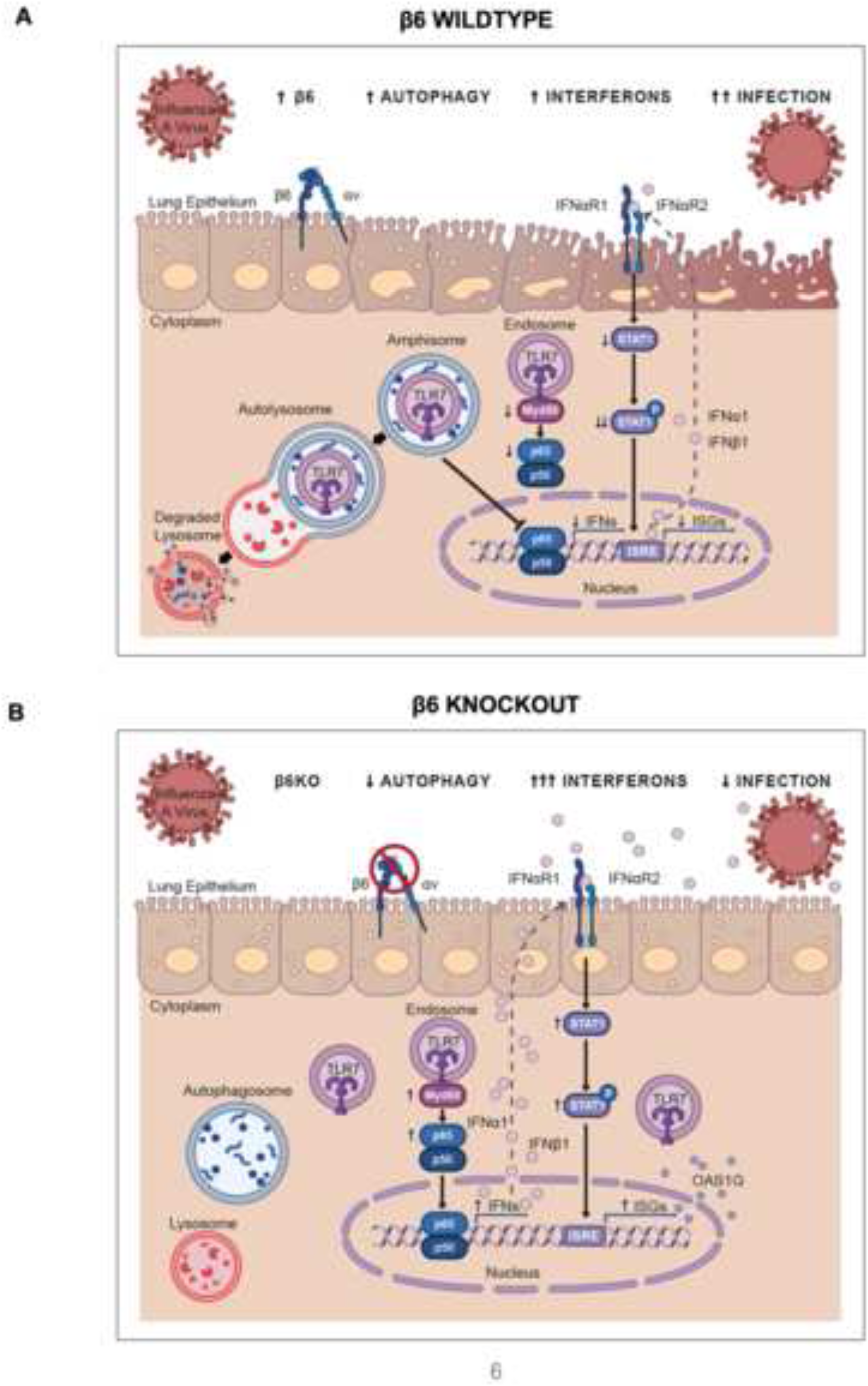
Model of β6 integrin regulation of epithelial-derived type I IFN responses during IAV infection. (A) Within the wildtype respiratory epithelium, IAV-induced β6 increases autophagy induction, resulting in the sequestration of endosomes within autophagosomes and increased trafficking of TLR7 sensors to acidic lysosomal compartments. This leads to a subdued TLR7-NFκB-pSTAT1 response against IAV infection. (B) The absence of the β6 integrin in the lung epithelium decreases the recruitment of autophagy machinery and reduces TLR7 trafficking to lysosomes, resulting in increased endosome availability and enhanced TLR7-mediated NFκB and STAT1 signaling against IAV infection.

We have provided evidence that the β6 integrin regulates TLR7 localization and signaling to modulate NFкB1 expression during IAV infection. Our findings support a similar role of integrins in regulating the TLR-mediated activation of NFкB1, as previously observed in macrophages (Han *et al*., 2010). This prolonged NFкB1 signaling is associated with sequential upregulation of IRFs in integrin-deficient cells, suggesting that integrins may also inhibit additional IFN-specific transcription factors along with NFкB1 (Han *et al*., 2010). Consistent with these studies, our RNA-seq and scRNA-seq analyses reveal significantly higher IRF7 and IRF9 levels in β6-deficient lung cells, suggesting an unidentified role of β6 as a transcriptional repressor of the epithelial immune response. Moreover, using the E6446 antagonist, we have demonstrated the dependency on TLR7 signaling to drive the NFкB1-pSTAT1 cascade in β6-deficient cells. However, it is important to note that E6446 also inhibits TLR9, which is targeted through integrin-mediated mechanisms as well (Lamphier *et al*., 2014; Acharya *et al*., 2016; Ueda *et al*., 2019). Given the shared endosomal localization and integrin regulation of these TLRs, future studies will explore if the role of β6 extends to the modulation of TLR9 localization, activation, and signaling in lung epithelial cells.

Aberrant signaling from TLR sensors has been implicated in the pathogenesis of various inflammatory and autoimmune diseases (Liew *et al*., 2005; Berland *et al*., 2006; Christensen *et al*., 2006; Marshak-Rothstein, 2006). Interestingly, despite the elevated type I IFN signaling dominating the β6KO lung microenvironment, we observed a significant reduction in inflammation, pro-inflammatory cytokine responses, and tissue damage compared to WT mice (Meliopoulos *et al*., 2016). In contrast, WT lungs exhibited significantly elevated TNF⍺ and IL6 levels, concomitant with pronounced alveolar permeability and inflammation (Meliopoulos *et al*., 2016). It is likely that this exuberant immune response observed in WT mice is due to overcompensatory immune mechanisms that attempt to supplement delayed IFN signaling in β6-expressing epithelial cells. As a result, to avoid detrimental inflammatory responses and maintain homeostasis, immune responses are tightly regulated by factors such as integrins (Lacy-Hulbert *et al*., 2007; Travis *et al*., 2007; Acharya *et al*., 2010). More recently, it has become clear that dysregulated TLR signaling is regulated by integrins since knockouts of ⍺v and ⍺v-associated β integrins upregulated the production of pro-inflammatory cytokines, elevated autoimmune antibodies, and led to persistent B lymphocyte activation (Han *et al*., 2010; Acharya *et al*., 2016, 2020). While these studies underscore the regulatory role of integrins in mediating TLR signaling in B-cells, the influence of β6-mediated TLR7 regulation on B-cell immunity remains to be determined (Soni *et al*., 2014). Identifying additional B-cell regulatory mechanisms will inform on the potential use of integrin agonists in mitigating the onset of autoimmune-related and inflammatory diseases.

Moreover, TLR ligands have been shown to trigger autophagy upon detection of pathogens (Delgado *et al*., 2008). It is possible that the low levels of autophagy observed in β6-deficient cells could promote the delivery of antigens to endosomal TLRs to facilitate pathogen elimination and further augment the antiviral response (Delgado *et al*., 2008). TLR-induced autophagy could also play a crucial role in averting IFN-triggered immunopathology through tight regulation of epithelial immunity in β6-deficient cells (Tian, Wang and Zhao, 2019).

The β6 integrin is involved in the activation of the transforming growth factor beta (TGFβ) by binding to the RGD motif on the TGFβ latency-associated peptide (John S. Munger *et al*., 1999; Annes *et al*., 2004; Giacomini *et al*., 2012). TGFβ is a pleiotropic cytokine with dichotomous roles, mediating both suppressive and inflammatory immune responses in a context-dependent manner (Sanjabi, Oh and Li, 2017). Previous studies have reported on the negative regulation of TLR-mediated signaling, NFкB1 activation, and associated IFN responses by TGFβ in natural killer cells, endothelial cells, and macrophages, respectively (Naiki *et al*., 2005; Eriksson *et al*., 2006; Lee *et al*., 2011; Hamon *et al*., 2022). Within the lung epithelium, TGFβ-deficient bronchial epithelial cells revealed a protective phenotype in IAV-infected mice, characterized by elevated IFNβ production and impaired viral replication (Denney *et al*., 2018). However, studies have not yet addressed the influence of the β6-TGFβ axis on immunomodulatory mechanisms within epithelial tissues. While our studies demonstrate the suppressive role of β6 on the IFN response, it will also be important to delineate if these mechanisms occur independently via β6 or through a TGFβ-dependent mechanism (Meliopoulos *et al*., 2016). Pharmacologically, current TGFβ inhibitors directly target receptor activity or cytokine function but due to its dual functionality and involvement in several biological processes, this systemic inhibition results in adverse effects on the host (Denney *et al*., 2018; Carvacho and Piesche, 2021; Essayas *et al*., 2023). By dissecting whether β6-activated TGFβ impairs intrinsic epithelial cell responses to infection, these studies may offer a more precise and localized approach to TGFβ inhibition for enhancing immunity while mitigating severe disease outcomes (Busenhart *et al*., 2022). Moreover, ⍺vβ1 also serves as a receptor for TGFβ, and in conjunction with ⍺vβ6, both are involved in the development of idiopathic pulmonary fibrosis (Decaris *et al*., 2021). Since inhibition of ⍺vβ1 and ⍺vβ6 have been shown to improve airway health, it would be interesting to assess whether there are any parallels between the ⍺vβ6-mediated inhibition of TLR7 signaling and the inhibitory effects of ⍺vβ1 on the immune response.

More recently, the association between integrins and autophagy components has been linked to TLR modulation (Acharya *et al*., 2016). Our proposed model complements these findings by demonstrating the integrin-mediated regulation of TLR7 signaling via autophagy induction. However, the mechanism by which integrins induce autophagy for immune modulation requires further investigation. Studies have implicated the Spleen tyrosine kinase (Syk) in driving intracellular integrin signaling to facilitate LC3 lipidation, resulting in the degradation of the Myd88 signaling adaptor or the colocalization of TLR sensors with LC3 machinery (Han *et al*., 2010; Acharya *et al*., 2016). Activated Syk suggests the induction of downstream mechanistic target of rapamycin (mTOR), thus implying the involvement of integrins in regulating mTOR to induce canonical macroautophagy (Leseux *et al*., 2006). However, preliminary data (not shown) suggest that the β6-activation of autophagy might not rely on mTOR regulation. It is possible that β6 may target other upstream autophagy regulators to induce selective autophagy (Zaffagnini and Martens, 2016). Alternatively, canonical and non-canonical TGFβ signaling pathways have also been shown to induce autophagy, evidenced by increased LC3 puncta formation, LC3B lipidation, and autophagosome-lysosome colocalization in epithelial cells (Trelford and Di Guglielmo, 2020, 2021). Considering the interconnectivity between β6 and TGFβ, this may shed light on whether β6 indirectly activates autophagy through TGFβ modulation. Furthermore, the specific mechanism by which epithelial-derived TGFβ attenuates the IFN response has yet to be elucidated. Exploring the intersection between TGFβ and autophagy pathways could offer valuable insights into this immunosuppressive mechanism.

The final stage of autophagy involves the lysosomal-induced degradation of delivered cargo, marking the completion of this process. Given that we observed TLR7-LAMP2a colocalization, we predict that this association may trigger the lysosomal-mediated degradation of this viral sensor, thereby terminating the TLR7 signal. As a result, we propose that treatment with lysosomal inhibitors, such as chloroquine and bafilomycin, should rescue the TLR7 signal in β6-expressing cells (Yamamoto *et al*., 1998; Yang *et al*., 2013; Mauthe *et al*., 2018; Fedele and Proud, 2020). However, it is important to note that these inhibitors impede autophagosome degradation through the neutralization of lysosomal and endosomal pH, which as a result inhibits TLR7 ligand binding and signaling (Macfarlane and Manzel, 1998; Yi *et al*., 1998; Kužnik *et al*., 2011). This caveat poses a challenge in testing this hypothesis. Chen *et al*. recently discovered a novel autophagy inhibitor, CUR5g, which blocks autophagosome-lysosome fusion without neutralizing intracellular vesicle acidity (Chen *et al*., 2022). Although the degradative effects of LAMP2a on TLR7 have not been demonstrated, the use of CUR5g could facilitate this investigation.

In summary, our data provide new evidence of the role of the β6 integrin in mediating TLR7 signaling in respiratory epithelial cells. These findings reveal an unappreciated mechanism of innate immune modulation by an epithelial-intrinsic integrin during influenza viral infection. Mechanistically, β6 suppresses epithelial-derived type I IFN responses by recruiting autophagy machinery and sequestering TLR7 sensors to lysosomal compartments, leading to downregulated TLR7-mediated activation of IFN signaling and induction mechanisms. A better understanding of the mechanisms that regulate epithelial immunity is critical for informing the development of therapeutic strategies that promote pathogen clearance while alleviating inflammatory and autoimmune diseases. Considering that pharmacologically developed β6 inhibitors are available and being tested in ongoing pre-clinical trials, we propose the β6 integrin as a suitable target for therapeutic intervention (John *et al*., 2020; Decaris *et al*., 2021; Sime and Jenkins, 2022). Transient inhibition of β6 signaling could potentially augment vaccination-induced immune responses against influenza infection, whereas the use of β6 agonists might serve to mitigate the development of detrimental inflammatory and autoimmune diseases.

## ACKNOWLEDGEMENTS

Confocal microscopy images were acquired using the facilities at the St. Jude Cell and Tissue Imaging Center, which is supported by SJCRH and NCI P30 CA021765. We appreciate the technical expertise, training, image analysis assistance, and scientific discussion provided by Dr. Aaron Taylor, George Campbell, and Dr. De Chen at the Cell and Tissue Imaging Center. We also thank Nathan Kurtz and Dr. Camenzind Robinson from the Electron Microscopy Shared Resource (SJCRH/ALSAC and NCI P30 CA021765), who provided assistance and expertise with the Transmission Electron Microscopy (TEM) images. Moreover, we gratefully acknowledge Dr. Shondra Miller, Mollie S. Prater, and Shaina Porter from the Center for Advanced Genome Engineering (NCI Grant P30CA021765) for generating the A549 β6KO cell line. We are extremely appreciative of Dr. Paul Thomas, Dr. David Boyd, and Dr. Jeremy Chase Crawford, including other involved members of the Thomas laboratory for the scRNA-Seq dataset. We also thank the St. Jude Hartwell Center for Biotechnology along with the Animal Resource Center and Veterinary Pathology Core that assisted with sequencing and animal handling, respectively. Many thanks to Marta Kosacheva for creating the graphical summary that was created using BioRender, Adobe Illustrator, and Adobe Photoshop.

This work was supported by ALSAC, the National Institute of Allergy and Infectious Diseases (NIAID) at the National Institutes of Health (NIH) under the Department of Health and Human Services (HHS) contract HHSN27220140006C for the St. Jude Center of Excellence for Influenza Research and Surveillance (CEIRS), by the NIAID contract 75N93021C00016 for the St. Jude Center for Excellence for Influenza Research and Response (CEIRR) to S.S-C. The St. Jude Graduate School of Biomedical Sciences (SJGS) provided support for M.S. The content is solely the responsibility of the authors and does not necessarily represent the views of NIH.

## AUTHOR CONTRIBUTIONS

**Conceptualization**: M.S., T.B., S.S.-C.; **Methodology**: M.S., T.B., V.M., B.S., P.H.B., J.C.C., M.S.P., S.M.P.-M., S.S.-C; **Data Curation**: M.S., T.B.; **Formal Analysis**: M.S., S.T., J.C.C., T.B.; **Investigation**: M.S., T.B., V.M., S.S.-C.; **Project Administration**: M.S., S.S.-C.; **Supervision**: V.M., J.C.C., S.S.-C.; **Writing – Original Draft**: M.S. **Writing – Revision and Edits**: M.S., T.B., V.M., P.H.B., S.T., J.C.C., S.S.-C.; **Funding Acquisition**: S.S.-C.

## STAR METHODS

### Key Resource Table

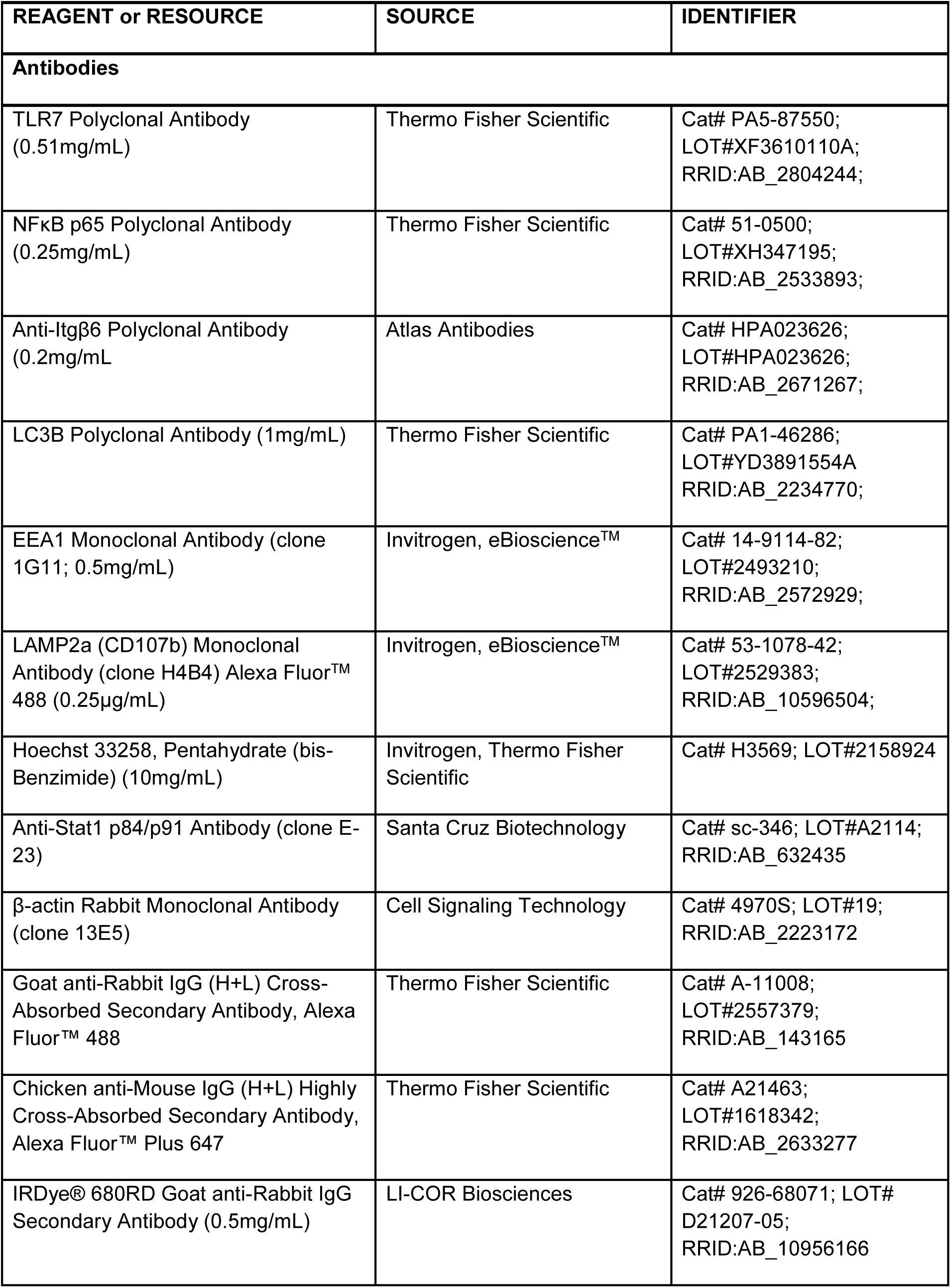
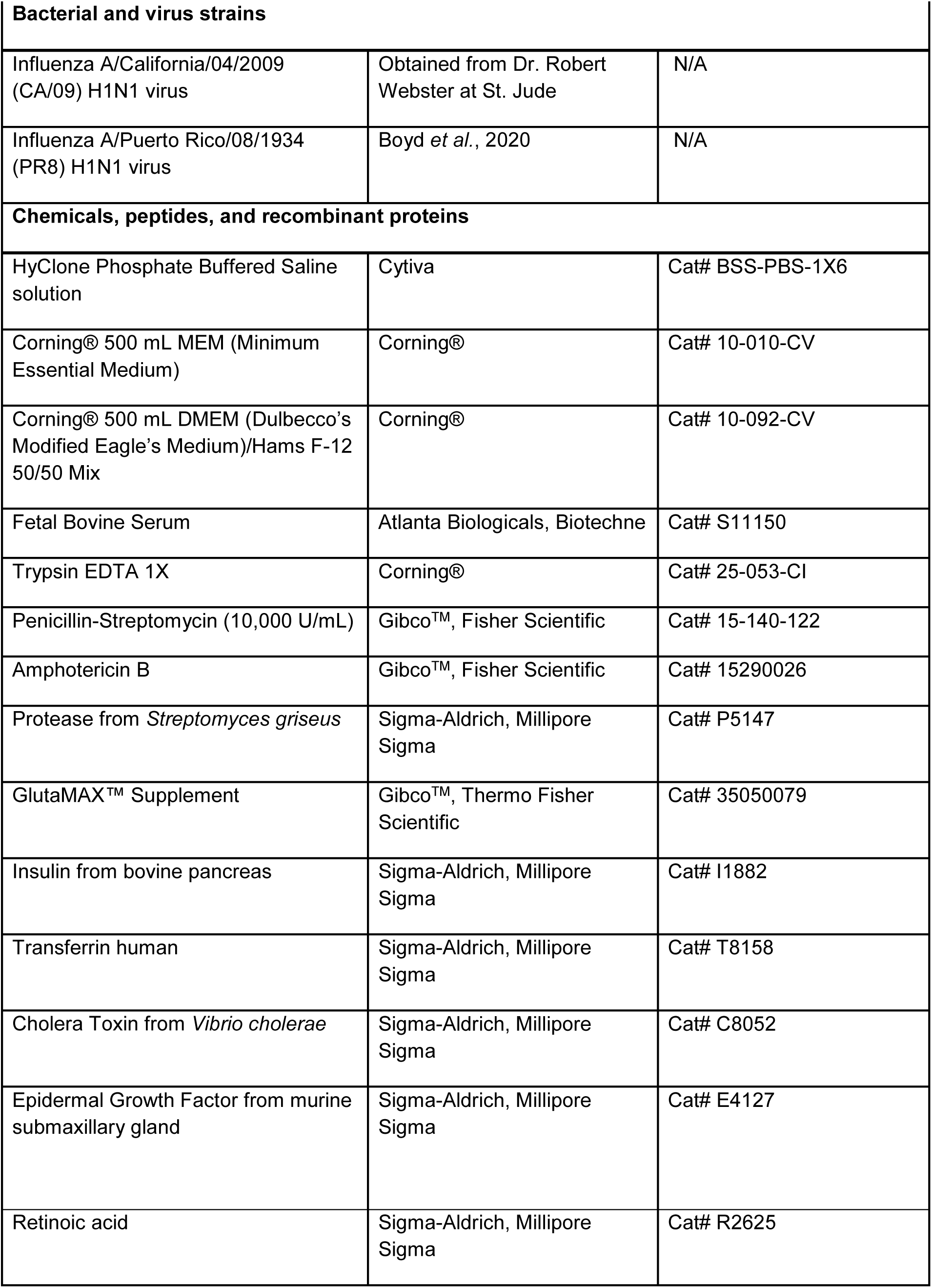
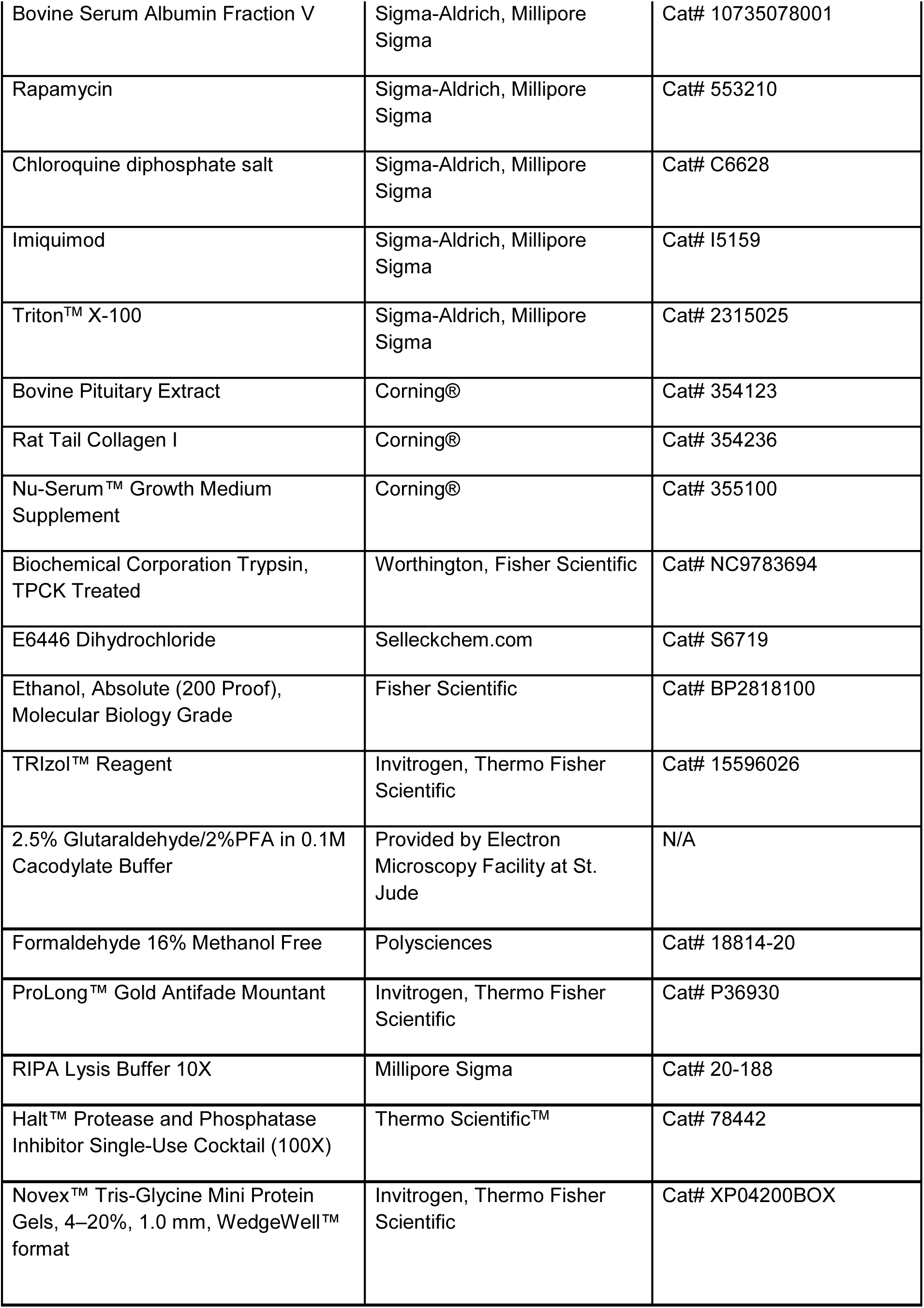
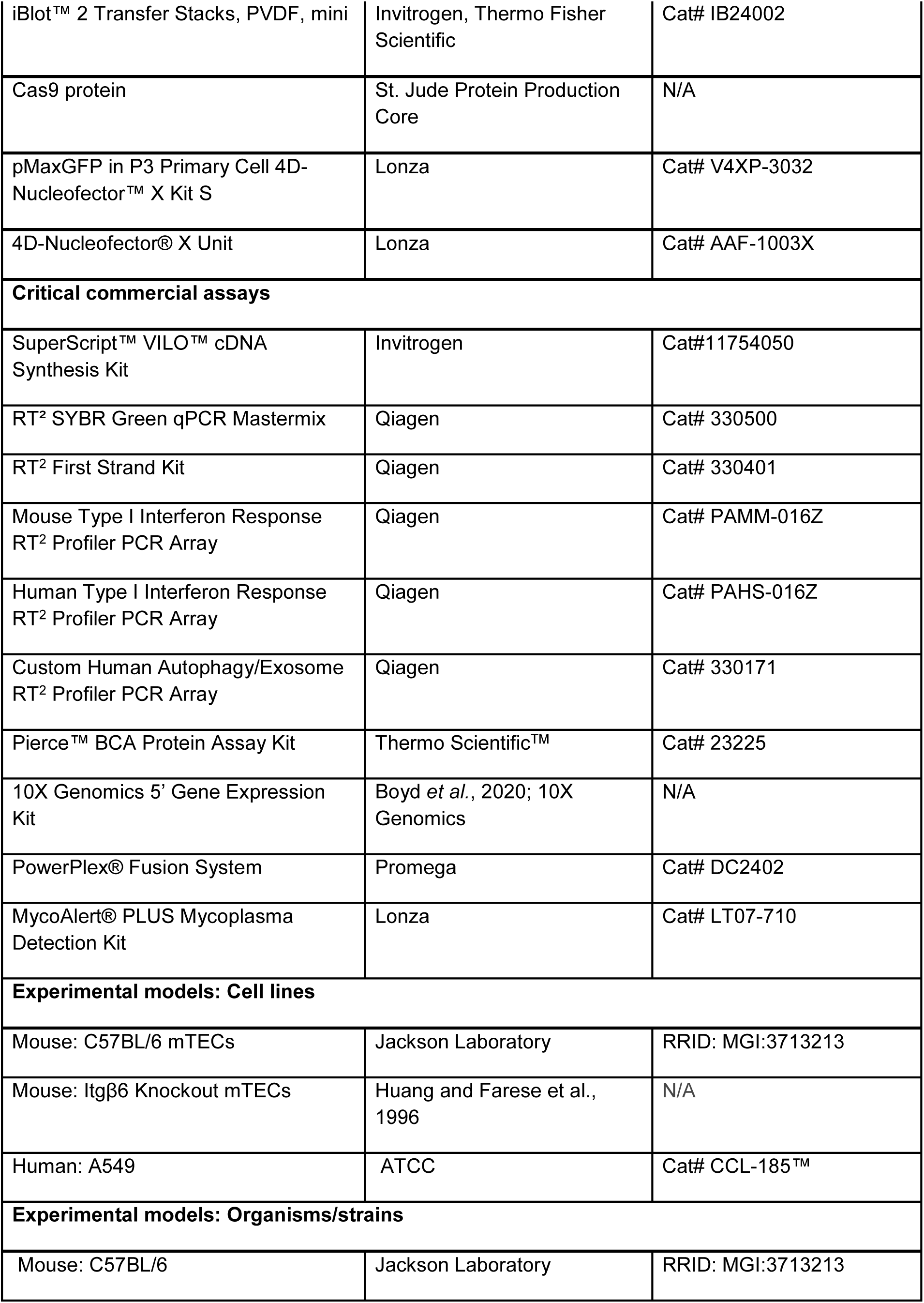
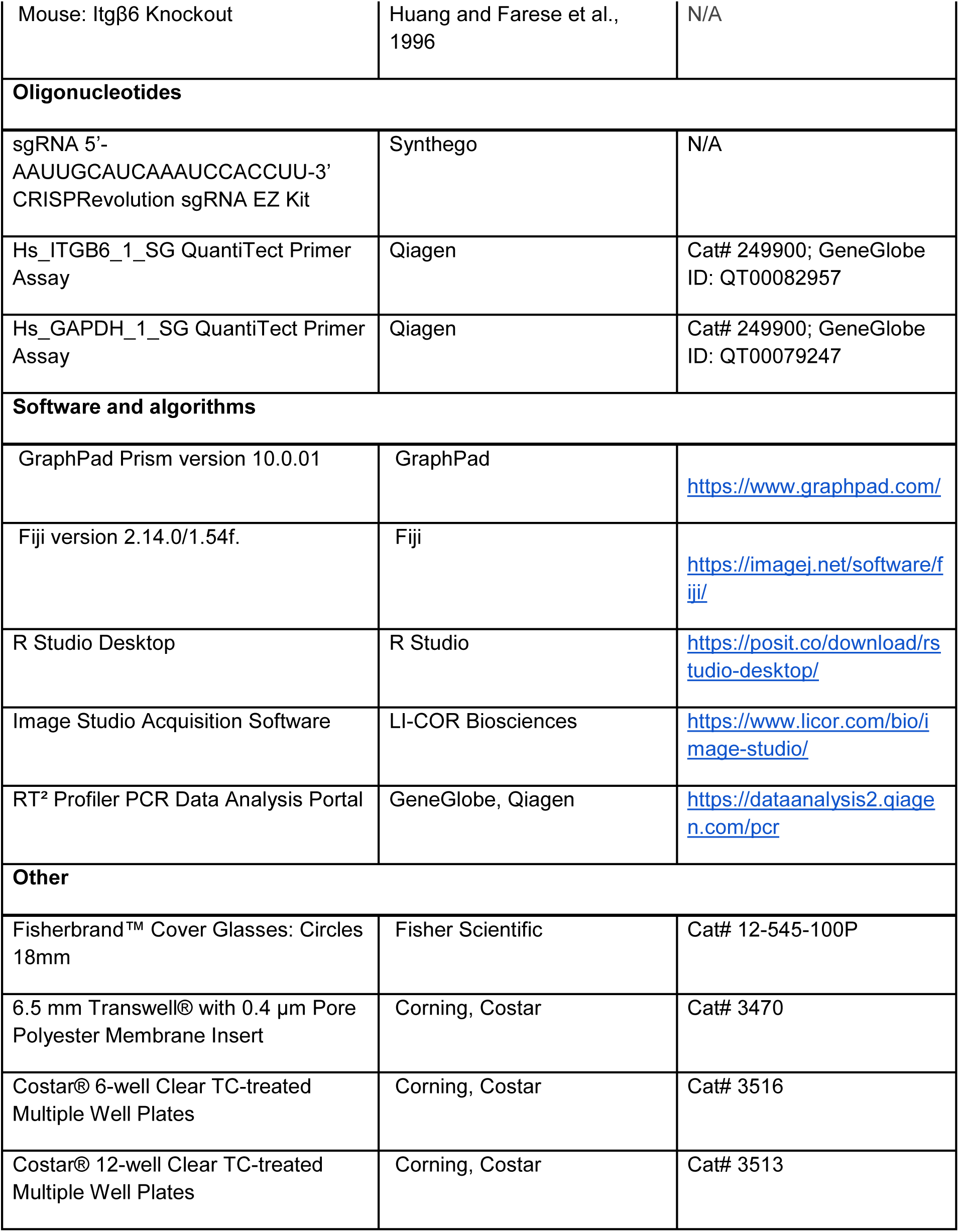

### Resource Availability

#### Lead Contact

Further information and requests for resources and reagents should be directed to and will be fulfilled by the Lead Contact, Dr. Stacey Schultz-Cherry.

#### Material Availability

- The A549 β6KO cell line generated for this study is available from the lead contact with a completed material transfer agreement.

#### Data and Code Availability

- This paper analyzes existing, publicly available RNA-sequencing data that has been deposited in the GEO database with accession number GSE68802 under https://doi.org/10.1371/journal.ppat.1005804.
- This paper analyzes existing, publicly available single-cell RNA sequencing data on the NCBI Short Read Archive under BioProjects PRJNA612345 under https://doi.org/10.1038/s41586-020-2877-5.

### Experimental Model and Subject Details

#### Mice

Eight- to ten-week-old C57BL/6 wildtype (WT) mice (Jackson Laboratory) were ordered from Jackson Laboratories. *Itgβ6*-/- mice were generated by the Dean Sheppard (UCSF) lab as previously described using mice from the C57Bl/6 background (Huang and Farese, 1996). All mice were bred in the Animal Resource Center at the St. Jude Children’s Research Hospital. All *in vivo* procedures were approved by the St. Jude Children’s Research Hospital Institutional Animal Care and Use Committee (IACUC) and followed in accordance with the Guide for the Care and Use of Laboratory Animals. These guidelines were established by the Institute of Laboratory Animal Resources and approved by the Governing Board of the U.S. National Research Council. Mice were housed in designated animal facilities where ambient temperature, humidity, food, water, and 12-hour light-dark cycles were regularly maintained.

#### Cell Lines and Primary Cell Cultures

Adenocarcinoma human alveolar basal epithelial (A549) cells were purchased commercially (ATCC) and maintained in-house. ITGβ6-/- (β6KO) A549 cells were generated via CRISPR-Cas9 knockout of the *Itgβ6* gene by the Center for Advanced Genome Engineering (CAGE) at the St. Jude Children’s Research Hospital. Briefly, 400,000 A549 cells were transiently transfected with pre-complexed ribonuclear proteins (RNPs) consisting of 100pmol of chemically modified sgRNA (5’ – AAUUGCAUCAAAUCCACCUU-3’), 35pmol of Cas9 protein, and 500ng of pMaxGFP via nucleofection according to the manufacturer’s recommended protocol using solution P3 and program CM-130 in a small 20µL cuvette. Five days post nucleofection, cells were single-cell sorted by FACs at the St. Jude Flow Cytometry and Cell Sorting Shared Resource to enrich for transfected, GFP-positive cells into 96-well tissue culture treated plates. Cells were clonally expanded and screened for the desired targeted modification (out-of-frame indels) via targeted deep sequencing using gene specific primers with partial Illumina adapter overhangs (CAGE1400.ITGB6.F – 5’ ACCTACATTTGGATTCAAGCACA −3’ and CAGE1400.ITGB6.R – 5’

GAAAGGACAGTCCCCATTTCAAC −3’, overhangs not shown) as previously described (Narina, Connelly and Pruett-Miller, 2023). NGS analysis of clones was performed using CRIS.py (Connelly and Pruett-Miller, 2019). Final clones were authenticated using the PowerPlex® Fusion System performed at the Hartwell Center for Biotechnology at the St. Jude Children’s Research Hospital. Final clones tested negative for mycoplasma by the MycoAlertTMPlus Mycoplasma Detection Kit. Knockout of the Itgβ6 gene from the A549 cell line was validated via RT-qPCR and confocal microscopy. No significant differences in cell morphology, doubling time, and response to various treatment conditions were observed between cell lines. Both cell lines were cultured in 1X MEM supplemented with 10% FBS and 100nM GlutaMAX.

Tracheas were harvested from euthanized mice, sliced longitudinally, and digested (DMEM:F12, 100µg/mL penicillin/streptomycin, 0.25µg/mL Amphotericin B and 0.15% pronase) overnight at 4°C. The overnight digest was inactivated with 10% FBS and the epithelial surface of the trachea was gently scraped into a 100cm dish containing Tracheal Epithelial Cell (TEC) media (DMEM:F12, 200nM GlutaMAX, 100µg/mL penicillin/streptomycin, 10µg/mL insulin, 5µg/mL transferrin, 100ng/mL cholera toxin, 125ng/mL epidermal growth factor, 15mg/500mL bovine pituitary extract, 5% fetal bovine serum and 5×10^-8^ retinoic acid). To isolate and eliminate fibroblasts, cells were incubated at 37°C in 5% CO_2_ for 3-4 hours to allow for fibroblast attachment. Supernatant-derived epithelial cells were seeded onto permeable transwell inserts pre-coated with 5µg/cm^2^ of rat-tail collagen at a density of 1×10^5^ cells/cm^2^. At complete confluency, the apical media was aspirated to establish the air-liquid interface (ALI) while the basolateral media was replaced with ALI media (DMEM:F12, 200nM GlutaMAX, 100µg/mL penicillin/streptomycin, 2%

NuSerum and 5×10^-8^M retinoic acid). Fully differentiated cells exhibited cilia formation and mucus production after 4-5 days. Cell medium was changed every 2 days.

#### Bacterial Cultures

This study did not use bacterial cultures.

### Method Details

#### Infections and Cell Treatments

Influenza A/California/04/2009 (CA/09) H1N1 virus was obtained from Dr. Robert Webster (St. Jude Children’s Research Hospital) and propagated in the allantoic cavity of 10-day-old pathogen-free embryonated chicken eggs. For *in vivo* infections, eight- to ten-week-old C57Bl/6 WT and β6KO mice were lightly anesthetized via isoflurane and then intranasally inoculated with 30µL CA/09 virus TCID50 10^3^ in PBS. Prior to the *ex vivo* infections, fully differentiated WT and β6KO mTECs were washed twice with 1X PBS for removal of excess mucus. CA/09 virus was diluted to a multiplicity of infection (MOI) of 0.01 in 1X MEM supplemented with 200mM GlutaMAX and 0.075% BSA. Apical transwell compartments were infected with 100µL viral inoculum per well and incubated at 37°C in 5% C0_2_ for 2 hours. Following viral absorption, the inoculum was aspirated and apical compartments were washed twice with 1X PBS. For *in vitro* infections, confluent monolayers were washed with 1X PBS. CA/09 virus was diluted to an MOI of 1 in infection media (1X MEM, 1% BSA, and 1% GlutaMAX). Washed monolayers were inoculated with 500µL/well CA/09 inoculum per well and incubated at 37°C in 5% C0_2_ for 1 hour. Following viral absorption, the inoculum was aspirated and replaced with infection media supplemented with 0.5µg/mL trypsin-TPCK. To inhibit TLR7 signaling, 10µM E6446 in 100% ethanol was diluted in infection media and used to pre-treat confluent A549 cells for 24 hours. To induce autophagy, A549 cells were treated for 24 hours with 100nM rapamycin diluted in infection media. To inhibit autophagosome degradation, infection media was supplemented with 8µM Chloroquine following one-hour viral absorption and incubated until sample collection. TLR7 positive control cells were incubated with 10µg/mL imiquimod diluted in infection media for 24 hours.

#### RNA Extractions, RT-qPCR and RT^2^ Profiler

PBS-washed cell monolayers (1.5×10^5^/250µL/well in 12-well plate) were treated with 750µL TRIzol reagent. Phenol-chloroform RNA extraction was performed on lung homogenates and cell lysates as described by the manufacturer. Eluted RNA was resuspended in 40µL nuclease-free water and RNA quality was assessed spectrophotometrically via Nanodrop. SuperScript VILO cDNA synthesis kit was utilized for cDNA synthesis. cDNA was diluted at 1:10 in nuclease-free water for quantitative PCR on the QuantiNova platform using QuatiTect primers. Human primers included *ITGβ6* (QT00082957) and *GAPDH* (QT00079247). A549 RNA lysates were diluted to 500µg and mTEC RNA lysates were diluted to 40µg. A549 and mTEC lysates were both reversed transcribed using the RT^2^ First Strand Kits. The cDNA was combined with the RT^2^ SYBR Green qPCR Mastermix and utilized in the following arrays - Mouse Type I Interferon Response, Human Type I Interferon Response, and Custom Human Autophagy/Exosome Array. Cycle threshold (CT) values were imported onto the Qiagen data analysis web portal (http://www.quiagen.com/geneglobe). The control group was assigned to the mTEC or A549 0 hpi sample while the test groups comprised β6KO 0hpi (all arrays) and A549 8 hpi and β6KO 8 hpi (custom array only). The data were normalized via manual selection of built-in reference genes (Actb, B2m, Gapdh, Hprt1, and Rplp0). Represented values are indicative of fold-regulation, which represents biologically relevant fold-change (2^-(Δ ΔCT)) data.

#### RNA-Sequencing

Fully differentiated WT and β6KO mTECs were washed twice with 1X PBS for removal of excess mucus. Trizol-extracted RNA was collected at baseline and samples were run in duplicate. 100ng RNA was converted to biotin-labeled cDNA following the Ambion protocol. The resulting cDNA was used by the Hartwell Center for Biotechnology at the St. Jude Children’s Research Hospital for hybridization to a Mouse Gene 2.0 array (Affymetrix). Quality assessment of raw sequencing data was performed using FastQC (https://www.bioinformatics.babraham.ac.uk/projects/fastqc/). Low-quality reads were trimmed with Trimmomatic version 0.39 (Bolger, Lohse and Usadel, 2014). Quality assessment was repeated once more with FastQC for confirmation of data quality improvements. Trimmed reads were aligned to the reference sheep genome (mm10 using STAR version 2.7.1a) and uniquely mapped reads were counted using feature Counts in the Subread package version 1.5.1 (Dobin *et al*., 2013; Liao, Smyth and Shi, 2014). Differential analysis was performed with the R package DESeq2 version 1.40.2 (Love, Huber and Anders, 2014).

#### Transmission Electron Microscopy

A549 and A549 β6KO cells were plated in a 6-well plate at a density of 1.5×10^5^/1mL per well. At ∼70% confluence, cells were infected at MOI 1 as described above. Positive control cells were treated with 100nM rapamycin for 24 hours. At 0 hpi, 8 hpi, 24 hpi, and 24-hour rapamycin treatment, cells were fixed in 2mL 2.5% Glutaraldehyde/2%PFA in 0.1M Cacodylate Buffer. Samples were subsequently fixed in osmium tetroxide and contrasted with aqueous uranyl acetate. Following this, samples were dehydrated using increasing concentrations of ethanol up to 100%, which was followed up with 100% propylene oxide. Samples were infiltrated with EmBed-812 and polymerized at 60°C. Embedded samples were sectioned at ∼70nm using a Leica ultramicrotome (Wetzlar, Germany) and examined with a ThermoFisher Scientific TF20 transmission electron microscope (Hillsboro, OR) at 80kV. Digital micrographs were captured via an Advanced Microscopy Techniques imaging system (Woburn, MA). All reagents were sourced from Electron Microscopy Sciences (Hatfield, PA), unless otherwise specified. For quantification analyses, an unbiased dataset of 63-120 images was collected from a total of n=2 cells per group for each experimental condition.

#### Single-cell RNA Sequencing Sample Preparation and Sequencing Analysis

The single-cell data was acquired from a dataset generated previously at St Jude (Boyd *et al*., 2020). Briefly, C57Bl/6 and β6KO mice (n=5 per group) were intranasally inoculated with mouse-adapted influenza A/Puerto Rico/08/1934 (PR8) virus at 2500 egg infectious dose 50 (EID50). Perfused lungs were digested, filtered, and subjected to red blood cell lysis. Cells were stained with viability and leukocyte markers, which allowed for the sorting of live, CD45-negative cells (Boyd *et al*., 2020). Sorted cells from each mouse were individually tagged with oligos and pooled for each time point. Approximately 25,000 cells per sample were loaded onto the 10X Genomics Chromium controller for partitioning of single cells into gel beads. The 10X Genomics 5’ Gene Expression Kit (version 2) was used to generate single-cell transcriptomic libraries, which were sequenced via Illumina NovaSeq (Boyd *et al*., 2020).

Initial processing of the 10X transcriptomics data was performed using CellRanger count (version 3.0.2, 10X Genomics). mm10 was used as the reference, which was altered to incorporate the PR8 genome. Individual samples were aggregated using Cell Ranger and normalized by the median number of mapped reads per identified cell. Normalized gene-expression matrices were then imported into Seurat (version 4.1.1) for downstream transcriptomic analyses and data visualization. Data were first filtered through the exclusion of any gene that was absent in at least 0.1% of total called cells. Cells that exhibited extremities in the total number of transcripts expressed (>30,000), the total number of genes expressed (<100 or >5500), or the percent of mitochondrial reads (>8%) were also excluded from downstream analyses. Data were normalized using default parameters that performed log transformation and variable features were identified via the vst method. Individual cell populations and sets were annotated using established markers specified in the literature (Boyd *et al*., 2020).

To assess the impact of influenza infection on the epithelial immune response, a subset of epithelial clusters was generated. The fastMNN algorithm was used for dataset integration from distinct libraries, which minimized subject- and sample-specific differences and allowed the identification of similar transcriptional subsets. The first 15 fastMNN dimensions were used for tSNE dimensionality reduction and nearest-neighbor graph construction to identify transcriptional clusters in Seurat. Detected epithelial cells were categorized into two groups: β6-positive (β6+) epithelial cells that expressed at least one Itgβ6 gene and β6-negative (β6-) epithelial cells where Itgβ6 expression was not detected. The AddModuleScore function in Seurat was used to calculate a comprehensive type I IFN response score based on a 19-gene signature consisting of well-established mediators of the type I IFN pathway: “*Nod1*”, “*Nod2*”, “*Casp1*”, “*Nlrp3*”, “*Aim2*”, “*Tlr3*”, “*Ticam1*”, “*Tlr7*”, “*Tlr9*”, “*Myd88*”, “*Ddx41*”, “*Trim56*”, “*Zbp1*”, “*Ripk3*”, “*Nfkb1*”, “*Irf7*”, “*Irf9*”, “*Tyk2*” and “*Stat1*”). The VlnPlot and DotPlot functions in Seurat were used to visualize gene expression differences between β6+ and β6-epithelial cells.

#### Immunofluorescence Microscopy

Autoclaved glass coverslips were added to the wells of a 12-well plate. A549 and A549 β6KO cells were seeded onto coverslips at 1.5×10^5^/500µL per well and incubated for 48 hours. Fully differentiated mTECs were seeded onto transwell inserts at 1×10^5^ cells/cm^2^/100µL per well and differentiated at the air-liquid interface as described above. At full confluence, monolayers were inoculated with CA/09 virus MOI 1 or treatment conditions as specified above. At designed collection time points, cells were fixed in 4% paraformaldehyde in PBS at room temperature for 20 minutes and washed twice with PBS. Monolayers were permeabilized with 0.1% Triton X in PBS at room temperature for 15 minutes. Cells were washed in PBS, then blocked at room temperature for 1 hour with 5% NGS for targets: TLR7, p65, Itgβ6, and LAMP2a and 5% BSA for targets: LC3-II and EEA1. Antibodies were diluted in 1% specified blocking buffer and cells were incubated with primary antibody solution overnight at 4°C. Primary antibodies included TLR7 (1:100), p65 (1:100), Itgβ6 (1:100), LC3II (1:100), EEA1 (1:50) and LAMP2a (1:250). The following day, cells were washed with PBS and incubated in the secondary antibody solution for 45 minutes in the dark at room temperature. Secondary antibodies included Alexa Fluor 488 goat anti-rabbit and Alexa Fluor 647 goat anti-mouse. Secondary antibodies were diluted to 1:200 in respective blocking buffers for targets: TLR7, Itgβ6, LC32II, and LAMP2a and diluted to 1:400 for p65. Each secondary antibody solution was supplemented with 1:1000 Hoechst. After final incubation, cells were washed with PBS and mounted on glass slides using Prolong Gold Antifade Mountant and cured for at least 24 hours in the dark. Slides were imaged on the Zeiss LSM 780 Observer.Z1 using a Plan Apochromat 63X/1.4 objective lens. Image acquisition was completed using Zen Black 2012 SP 5 (14.0.28.201). Colocalization quantification was measured with Mander’s co-efficient using Fiji macro code generated by the Cell and Tissue Imaging Center at St. Jude.

#### Immunoblotting

A549 and A549 β6KO cells were plated in a 12-well plate at a density of 1.5×10^5^/500µL per well. After 48 hours post-plating, confluent monolayers were washed with cold PBS and incubated with either viral inoculum or treatment condition, as detailed above. At the designated time point, cells were collected with 100µL cold RIPA Lysis Buffer that was supplemented with 1X Halt protease inhibitor cocktail. The plates were incubated on ice for 2 minutes. Cell monolayers were scraped off, transferred to cold Eppendorf tubes, vortexed, and placed in ice for 30 minutes. Tubes were vortexed at 10-minute increments and then vortexed at 11,000g for 15 minutes at 4°C. Protein concentrations were quantified using the BCA Protein Assay Kit. Equivalent protein concentrations (50µg) were prepared under reducing conditions and loaded onto a 4-20% Tris-Glycine SDS-PAGE 1.0mm Mini Protein Gels. Gels were transferred to PVDF membranes using the iBlot 2 transfer stacks. Membranes were blocked in 1X TBS for 1 hour at room temperature and then incubated with the primary antibodies in 5% BSA/TBST at 4°C overnight. Primary antibodies included STAT1 at 1:1000 dilution and β-actin at 1:2000 dilution. The IRDye 680RD goat anti-rabbit IgG secondary antibody was used at 1:1000 dilution for 1-hour incubation at room temperature. Immunoblots were imaged using Image Studio Acquisition Software.

### Quantification and Statistical Analysis

All statistical analyses were performed with GraphPad Prism software version 10.0.01. For all experiments, information on biological and technical replicates, statistical tests, and p-values are reported in figure legends. Briefly, p-values less than 0.05 were considered to be statistically significant. Statistical significance was determined by performing a t-test for two unpaired groups or two-way ANOVA with Sidak’s multiple comparisons test for experiments that have two factors with multiple comparisons. Microscopy images were analyzed with Fiji version 2.14.0/1.54f.

---

**Figure S1.**
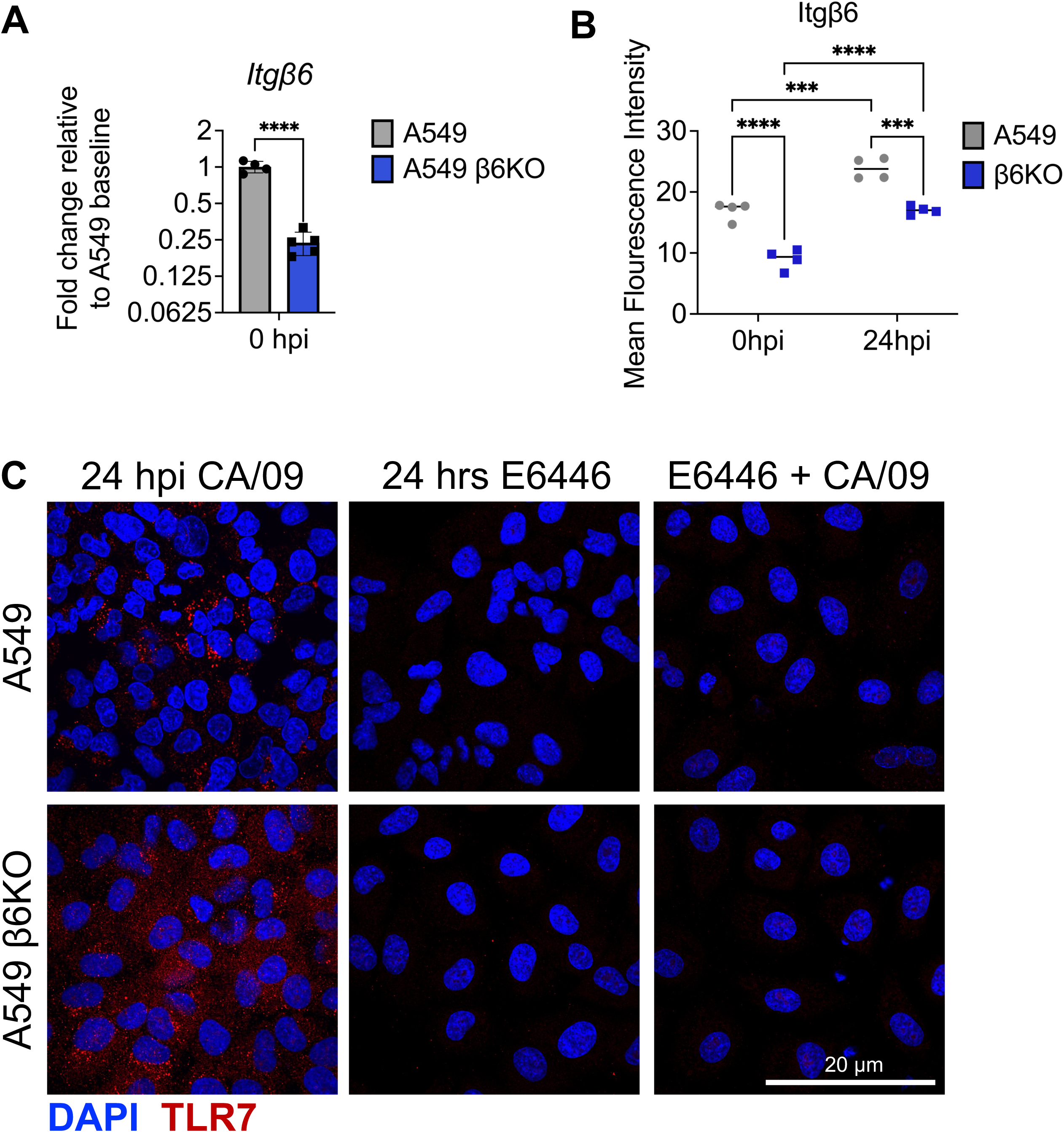
Validation of A549 β6KO cell line and E6446 inhibitor, Related to Figures 1-2 (A) RT-qPCR analysis of *Itgβ6* expression RNA isolated from A549 and A549 β6KO cells at baseline representative of 2 independent experiments with n=4-5 cells per group. (B) Mean fluorescence intensity of β6 expression in A549 and A549 β6KO cells at 0 hpi compared to 24 hpi. Data is from pooled 2 independent experiments with n=2 cells per group. (C) A549 and A549 β6KO cells were immunostained for TLR7 (red) and nucleus (Hoechst, blue). Data representative of 2 independent experiments with n=3 cells per group. First column to the left consists of cells infected with CA/09 virus MOI 1 at 24 hpi. Middle column consists of cells treated with 10µM E6446 TLR7 inhibitor for 24 hours. Final column to the right consists of cells pre-treated with 10µM E6446 and then infected with CA/09 virus MOI 1 for 24 hours. Scale bar, 20µm. Mean fluorescence intensity quantified with Fiji. Data are represented as mean ± SD. Unpaired T-test for A; Two-way ANOVA with Sidak’s multiple comparisons test for B.

**Figure S2.**
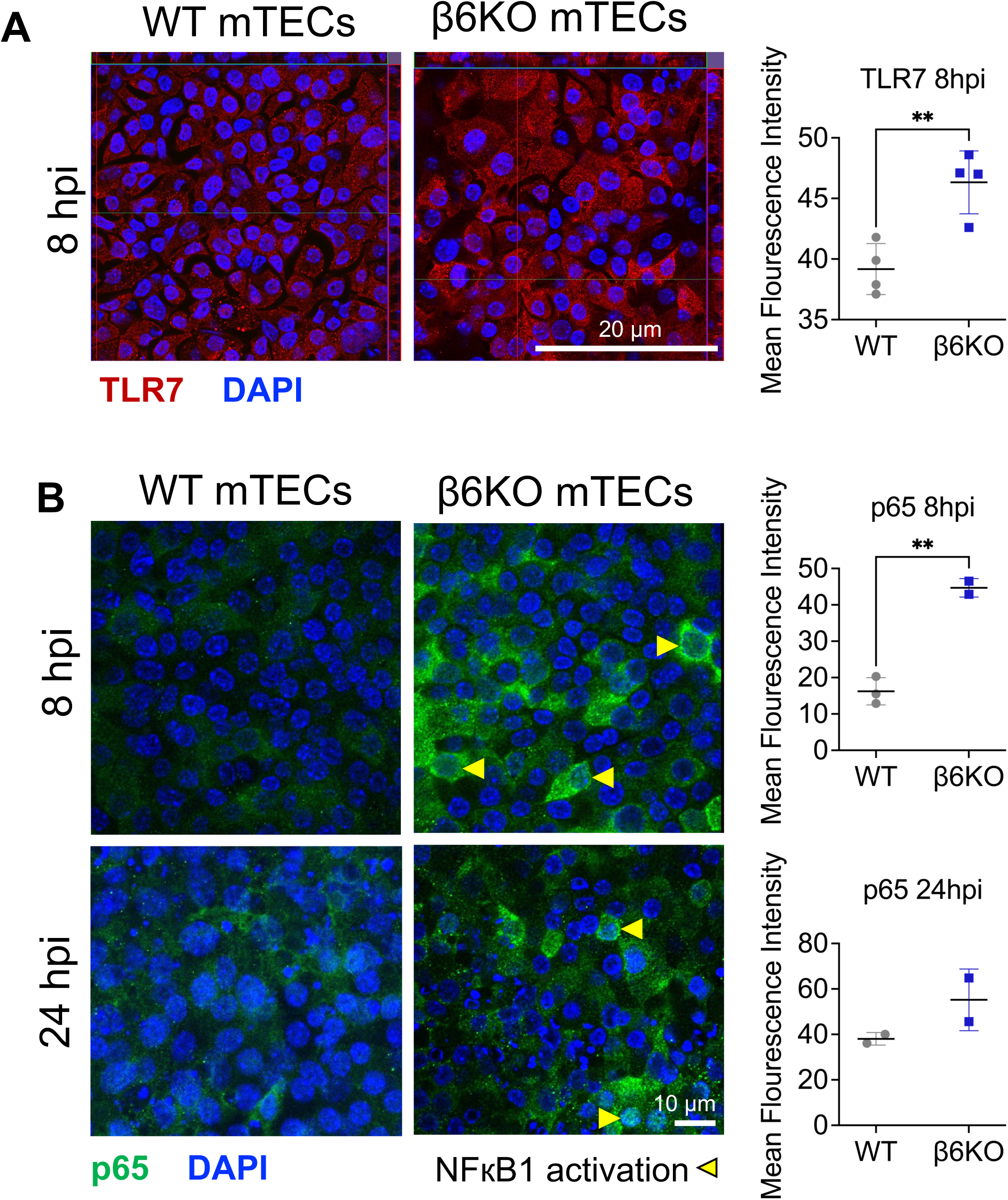
TLR7 and NFкB1 expression in IAV-infected mTECs, Related to Figure 2 (A) WT and β6KO mTECs were infected with CA/09 virus MOI 1 for 8 hpi and then immunostained for TLR7 (red) and nucleus (Hoechst, blue). Data pooled from 2 independent experiments with n=2 cells imaged. (B) WT and β6KO mTECs were infected with CA/09 virus MOI 1 for 8 and 24 hpi, then immunostained for NFκB subunit p65 (green) and nucleus (Hoechst, blue). Data representative of 2 independent experiments with n=2-3 cells per group. Yellow arrows indicate p65 nuclear localization. Scale bar, 10µm. Mean fluorescence intensity quantified using Fiji. Data graphed as mean ± SD via unpaired T-test for A and B.

**Figure S3.**
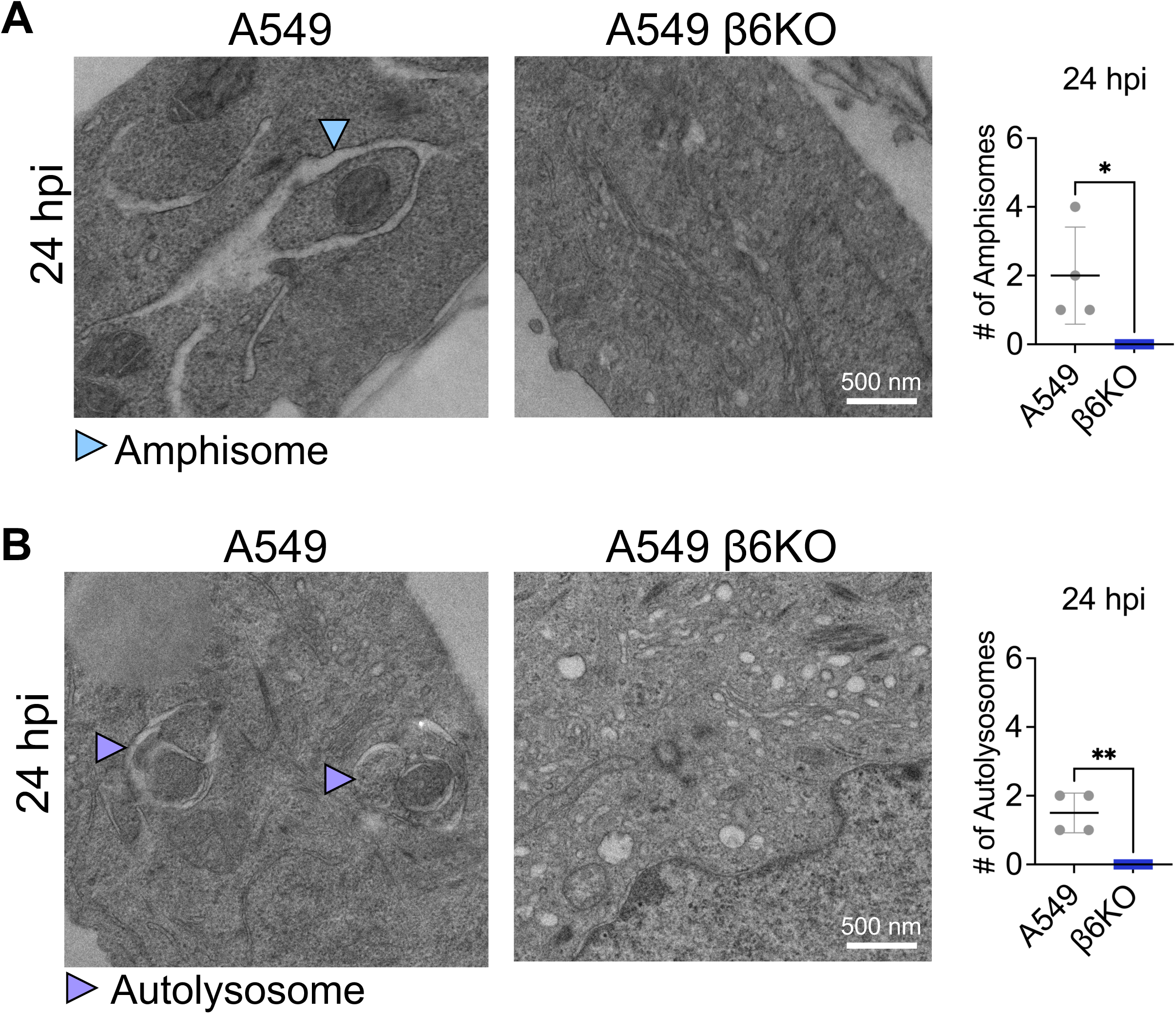
Additional autophagy machinery identified in IAV-infected A549s, Related to Figure 3 (A) TEM of amphisomes in A549 and A549 β6KO cells infected with CA/09 virus MOI 1 for 24 hpi. Blue arrow indicates amphisome. (B) TEM of autolysosomes in A549 and A549 β6KO cells infected with CA/09 virus MOI 1 for 24 hpi. Purple arrows indicate autolysosomes. For both amphisomes and autolysosomes, data were pooled from 2 independent experiments with n=1 cell per group for each treatment condition and quantified from 63-120 unbiased images. Scale bar, 500nm. Data graphed as mean ± SD via unpaired T-test. *p<0.05; ***p≤0.0001; ****p<0.0001.

**Figure S4:**
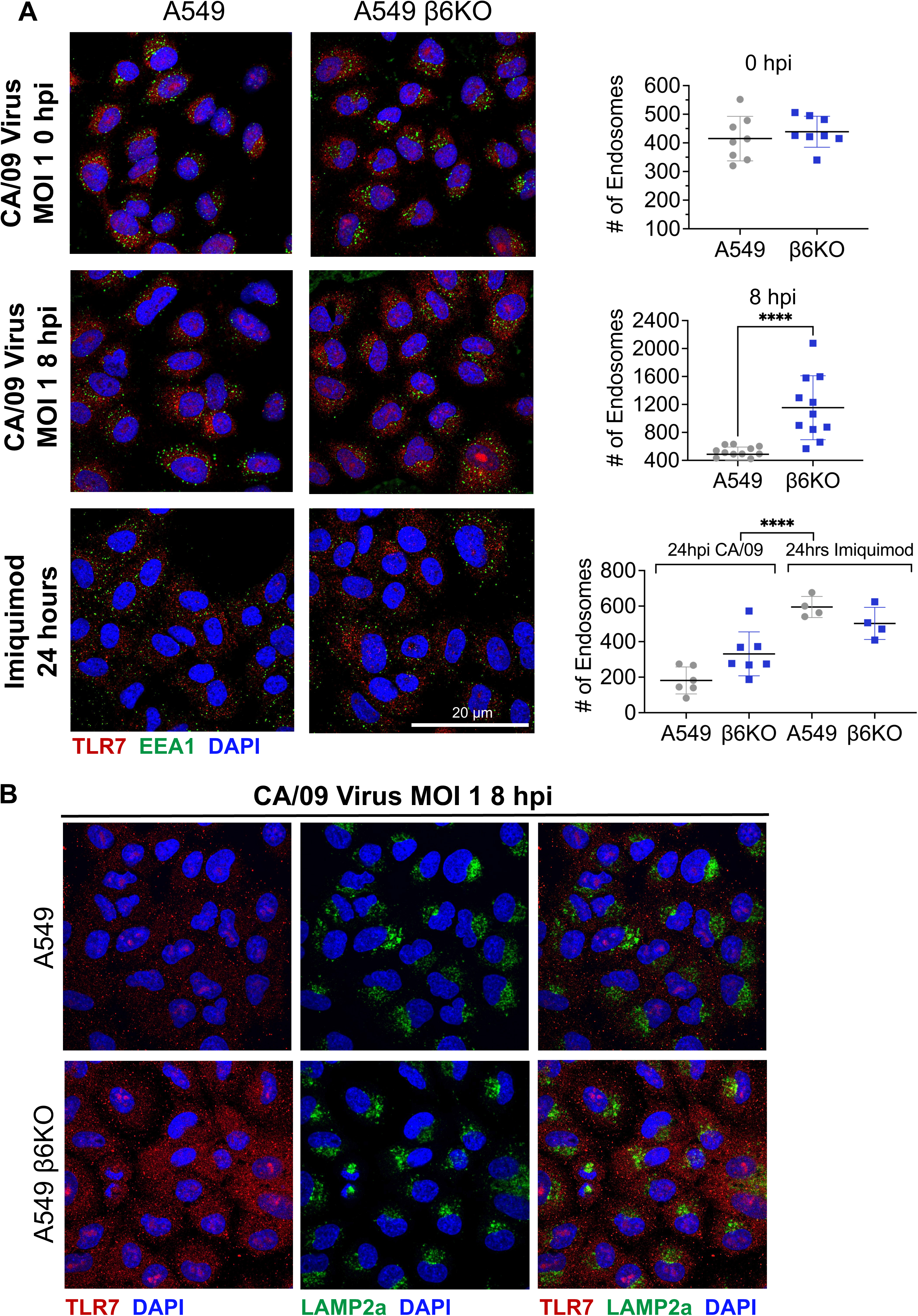
Comparison of endosome frequencies and lysosome expression in A549 cells at 8hpi, Related to Figure 4 (A) A549 and A549 β6KO were immunostained for TLR7 (red), EEA1 endosome (green), and nucleus (Hoechst, blue). Data pooled from 2 independent experiments with n=4-7 cells per group. First panel represents uninfected cells. Second panel indicates cells that were infected with CA/09 virus MOI 1 for 8 hours. Third panel represents cells that were incubated with 10µg/mL imiquimod (TLR7 agonist) for 24 hours. Number of endosomes quantified using Fiji. Scale bar, 20µm. (B) A549 and A549 β6KO were infected for 8 hours with CA/09 virus MOI 1 and immunostained for TLR7 (red), LAMP2a lysosome (green), and nucleus (Hoechst, blue). Data representative of 2 independent experiments with n=2 cells per group. Scale bar, 20µm. Data graphed as mean ± SD. Unpaired T-test for A (0 and 8hpi). Two-way ANOVA with Sidak’s multiple comparisons test for A (imiquimod 24 hours). ****p<0.0001.

